# Do Domain-Specific Protein Language Models Outperform General Models on Immunology-Related Tasks?

**DOI:** 10.1101/2023.10.17.562795

**Authors:** Nicolas Deutschmann, Aurelien Pelissier, Anna Weber, Shuaijun Gao, Jasmina Bogojeska, María Rodríguez Martínez

**Author notes:** A. Pelissier contributed equally with N. Deutschmann.

## Abstract

Deciphering the antigen recognition capabilities by T cell and B cell receptors (antibodies) is essential for advancing our understanding of adaptive immune system responses. In recent years, the development of protein language models (PLMs) has facilitated the development of bioinformatic pipelines where complex amino acid sequences are transformed into vectorized embeddings, which are then applied to a range of downstream analytical tasks. With their success, we have witnessed the emergence of domain-specific PLMs tailored to specific proteins, such as immune receptors. Domain-specific models are often assumed to possess enhanced representation capabilities for targeted applications, however, this assumption has not been thoroughly evaluated. In this manuscript, we assess the efficacy of both generalist and domain-specific transformer-based embeddings in characterizing B and T cell receptors. Specifically, we assess the accuracy of models that leverage these embeddings to predict antigen specificity and elucidate the evolutionary changes that B cells undergo during an immune response. We demonstrate that the prevailing notion of domain-specific models outperforming general models requires a more nuanced examination. We also observe remarkable differences between generalist and domain-specific PLMs, not only in terms of performance but also in the manner they encode information. Finally, we observe that the choice of the size and the embedding layer in PLMs are essential model hyperparameters in different tasks. Overall, our analyzes reveal the promising potential of PLMs in modeling protein function while providing insights into their information-handling capabilities. We also discuss the crucial factors that should be taken into account when selecting a PLM tailored to a particular task.

## 1 Introduction

T cells and B cells are integral to the adaptive immune system, each executing critical functions to orchestrate robust defenses against invading pathogens and other internal challenges [1]. Both types of cells are pivotal for both acquired immunity and antigen-specific responses due to their unique capability to recognize and respond to specific antigens and epitopes. Namely, T cells are responsible for surveillance and cytotoxic activities against infected or aberrant cells, while B cells undergo affinity maturation and can produce highly specific antibodies (Abs) to neutralize antigens. Both cells are essential for allowing the immune system to distinguish between self and non-self entities. Regarding recognition, both T and B cells detect foreign antigens via hyper-variable B cell and T cell receptors (BCRs and TCRs), however, the mechanisms of recognition differ. B cells identify free, unprocessed antigens, while T cells detect antigens presented by the major histocompatibility complex (MHC), a complex of cell surface proteins expressed on the surface of antigen-presenting cells.

At the molecular level, BCRs and TCRs are sequences of amino acids that form complex 3D structures that can undergo conformational alterations. Adaptive immunity somatically produces extensive repertoires of TCRs and BCRs, potentially enabling the recognition of many different non-self molecules. During their development, each T and B cell assembles a unique receptor through the rearrangement of different V, D, and J gene segments. This process is accompanied by random nucleotide insertions and deletions at the *junctions* between these gene segments, specifically at the V-D and D-J boundaries [2]. The unique combination of segments and their junctions forms the complementary determining region 3 (CDR3), which is the most diverse part of the sequence. This region governs the binding specificity of B cell-derived immunoglobulins and T cell receptors, thus influencing their downstream functions [3]. Due to the stochastic nature of the recombination process, a large diversity of receptors can be generated, each one with unique antigen specificity. Estimates of the theoretical diversity vary, but it can be as large as 10^20^ [4, 5]. While the theoretical diversity is huge, a smaller number of unique TCRs and BCRs is typically expressed in an individual, with estimates suggesting 10^13^ unique proteins in the human antibody repertoire [6]. This high diversity allows the adaptive immune system to respond to the myriad of both seen and unseen threats.

The recent advent of high-throughput adaptive immune receptor repertoire (AIRR) sequencing has provided unprecedented insights into the diversity and adaptability of our immune system, but also raised significant challenges in terms of data analysis and interpretation. While many predictive algorithms for antibody-antigen and TCR-epitope interactions have been developed, most exhibit suboptimal accuracy. This limits the rational design of antibodies [7], novel T-cell mediated therapies [8], and new forms of immunotherapy beyond oncological applications [9].

In recent years, self-supervised protein language models (PLMs) have emerged as a powerful paradigm for a large number of protein-related tasks, including biological and molecular property prediction [10]. At their core, these models treat amino acid sequences as a biological language. This language can be decoded using deep learning models trained on vast numbers of protein sequences (approximately 250 million), enabling the translation of specific sequences into meaningful vector representations of proteins (embeddings) in a high-dimensional latent space. Some of the best-known PLMs include protBERT (Bidirectional Encoder Representations from Transformers) [11], ESM (Evolutionary Scale Modeling) [12], aminoBERT [13] and ProGen [14]. Although neither of them was specifically trained for molecular property or structure prediction, their immense scale (15 billion parameters for the largest) allows them to distill fundamental qualities of the biological language. This is demonstrated by their ability to predict protein 3D-structures [15, 16], binding events [17], and identifying functional sites [18]. Importantly, the latent space representations generated by these models can be used as input for subsequent predictive models, reducing training time and model complexity, and boosting performance on downstream tasks. However, the size and complexity of these models raise substantial computational challenges, including the necessity of significant processing power and extensive training times. For instance, the largest among them, ESM2-15B, cannot operate on standard commercial machines and requires the use of costly clusters for embedding protein sequences. Therefore, when selecting a PLM, it is crucial to assess the model’s size in relation to expected performance in order to minimize computational costs while maintaining good accuracy.

### Immune-specific PLMs

Given the success of these PLMs in many bioinformatics-related tasks, similar models have been trained on immune-specific protein sequences, such as antibodies (AbLang [19], Antibert [20], AbMAP [21]) and T-cell receptors (TCR-BERT [22], catELMO [23]). The rationale behind these models is that they might improve representation capabilities for immune-related applications [24]. Nevertheless, as they are usually trained on a narrower and less diverse dataset of proteins, there is a risk of missing general knowledge that might help the model learn and generalize better for unseen proteins and antigens [21].

In this article, we explore diverse use cases of PLMs (Table 1) for immune-related tasks. Specifically, we use embeddings directly derived from PLMs as input for basic multi-layer perceptrons trained for antigen and epitope-related tasks. We also discuss the implications of employing general models over specialized immune-specific PLMs. Our objective is not solely to ascertain the peak performance on specific tasks but to also comprehensively discuss the crucial factors that should be taken into account when selecting a protein embedding model tailored to a particular task. In this context, we demonstrate that the prevailing notion of “domain-specific models outperform general models” requires a more nuanced examination. Additionally, we highlight the critical importance of selecting the appropriate layer for extracting embeddings in optimizing performance on downstream tasks.

**Table 1:** Characteristics of PLMs used to embed immune sequences in this article. All layers of a given PLM have the same dimension.

|  | ESM2 [15] |  |  |  | AbLang [19] | TCR-BERT [22] |
| --- | --- | --- | --- | --- | --- | --- |
| Input | Any protein | | | | BCR / Antibody | T-cell CDR3-β |
| Trained on | UniRef [25] |  |  |  | OAS [26] | PIRD[27], VDJdb[28] |
| Training set size | 60M |  |  |  | 14M | 100K |
| Nr. Parameters | 8M | 35M | 150M | 650M | ~250M | ~250M |
| Nr. Layers | 6 | 12 | 30 | 33 | 12 | 12 |
| Attention Heads | 20 | 20 | 20 | 20 | 12 | 12 |
| Embedding dim. | 320 | 480 | 640 | 1280 | 768 | 768 |

## 2 Results

### 2.1 The choice of layer as embedding from PLMs matters

We began by examining the similarities and differences between embeddings-based metrics and other established distances for sequence analysis such as the Levenshtein distance. Levenshtein distance commonly referred to as the *edit distance*, quantifies sequence dissimilarity by determining the minimal number of single-character edits needed to transform one string into another [29].

We calculated pairwise distances between sequences by measuring the Euclidean distances within various embeddings, and compared them to the normalized Levenshtein pairwise distances (Methods 4.2) using the Pearson and Kendal correlations (Fig. 1). The Levenshtein distance is a measure of the similarity between two strings that can be computed both at the nucleotide or amino acid level [30]. To be consistent with PLMs that operate at the amino acid level, in this paper we always computed it at the amino acid level. While both Ablang and ESM-650M showed a significant correlation with the Levenshtein distance, there were also substantial differences, with ESM2-650M obtaining higher Pearson coefficients than Ablang (0.76 and 0.57, respectively, Figure 1A & B). Interestingly, the pattern was inverted when we examined the CDR3-*β* of TCRs, with Pearson correlation coefficients of 0.73 and 0.49 for TCR-BERT and ESM2-650M, respectively (Figure S3).

**Figure 1:**
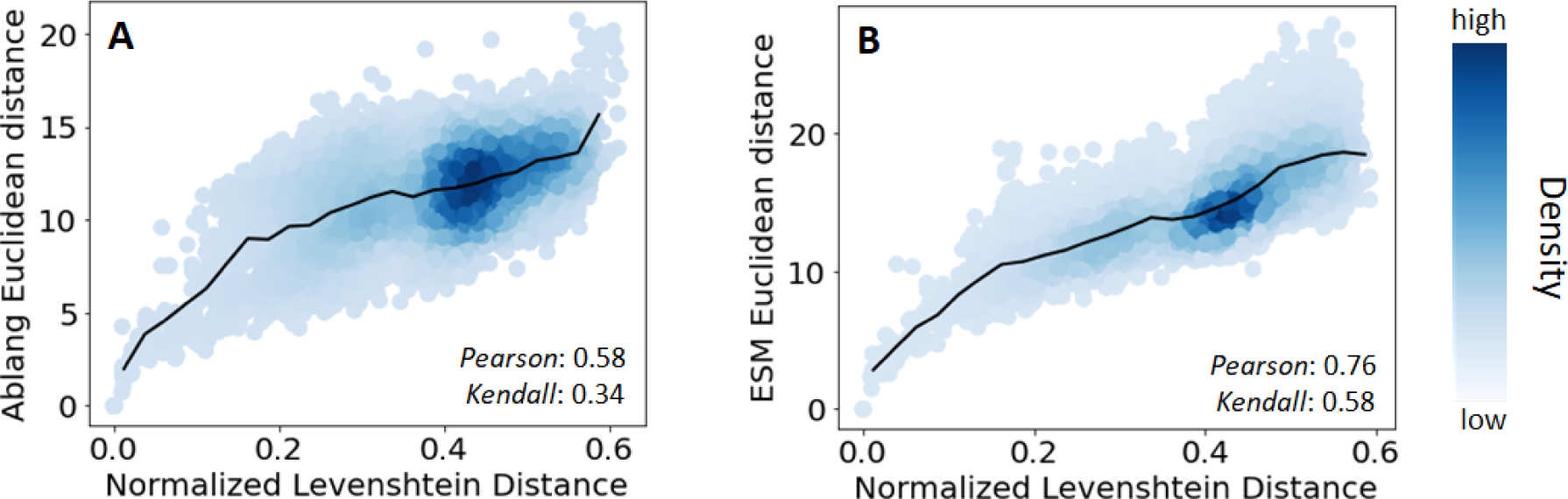
Relationship between Levenshtein and Euclidean distance in the (A) AbLang and (B) ESM2-650M-L6 embedding space. The sequences are BCR heavy chains taken from 10 germinal center B cells in the same human lymph node [31].

Before moving forward, we make two clarifications. First, for the TCRs, our comparison focused only on CDR3-*β* chains since TCR-BERT cannot encode full sequences. Second, a comparison with the Levenshtein distance serves at best as an indirect measure of representation quality, as a single amino acid difference minimally changes the Levenshtein distance between two sequences, but can drastically alter the receptor’s functional properties. In the next sections, we will provide more meaningful evaluations focusing on affinity prediction tasks.

Several studies on PLMs outside of biology have already emphasized the importance of selecting the appropriate layer for embedding extraction as one of the model’s hyperparameters [32, 33, 34]. Therefore, we swept across different layers of PLMs and evaluated the changes in correlation to the Levenshtein distance. Denoting the layer from which the embedding was taken as L*X*, we observed significant differences for both B cell and T cell receptors (Table 2). For example, using the L6 instead of L33 of ESM2-650M increased the Pearson correlation to 0.76 from 0.63 on BCRs and 0.59 from 0.49 on TCRs CDR3-*β*. Similarly, L8 of TCR-BERT yielded a higher correlation with the Levenshtein distance compared to L12. As we were solely evaluating the correlation with the Levenshtein distance, we do not claim that earlier layers are inherently more expressive than deeper ones. Our main observation is that the choice of layer should be considered as a hyperparameter that requires optimization for each specific task and embedding.

**Table 2:** Relationship of distance in various embeddings to Levenshtein distance of B cell receptors and T cell receptors sequences. Note that for compatibility with TCR-BERT, only CDR3-*β* sequences are considered for T cells.

|  | Pearson<br>(B cells) | Kendall<br>(B cells) | Pearson<br>(T cells) | Kendall<br>(T cells) |
| --- | --- | --- | --- | --- |
| AbLang-L12 | 0.58 | 0.34 | — | — |
| TCR-BERT-L8 | — | — | **0.75** | **0.52** |
| TCR-BERT-L12 | — | — | 0.73 | 0.50 |
| ESM2-650M-L6 | **0.76** | **0.58** | 0.59 | 0.39 |
| ESM2-650M-L33 | 0.63 | 0.41 | 0.49 | 0.32 |
| ESM2-150M-L30 | 0.63 | 0.42 | 0.44 | 0.23 |
| ESM2-35M-L12 | 0.60 | 0.47 | 0.42 | 0.30 |
| ESM2-8M-L6 | 0.40 | 0.35 | 0.44 | 0.30 |

### 2.2 Generalist and domain-specific PLMs for the characterization of B cells evolutionary trajectories

In this section, we explored the use of PLMs to characterize B cell evolution during adaptive immune responses, and evaluated the evolutionary information captured by the different embeddings. Briefly, upon infection, B cells undergo rapid proliferation within the germinal centers (GCs), where they mutate their BCR genes to optimize the binding affinity to the invading pathogen. Hence, GCs are critical BCR evolutionary hubs that are shaped by intricate selection dynamics and inter-GC communication events. As a result, tightly intertwined BCR clones co-evolving together and potentially shared across GCs are often found in experimental studies.

Typical B cell repertoire analyzes start by grouping BCR sequences into clones descending from a common ancestor using the Levenshtein distance. Next, phylogenetic lineages of clonally related B cells are reconstructed to elucidate evolutionary trajectories, identify clonal interrelationships, and investigate dynamic immune responses over time. Such analyzes can shed light not only on the emergence of functional antibodies but also on the progression of conditions such as chronic infections, autoimmune disorders, and cancer.

However, these analyzes rely exclusively on multiple sequence alignments and quantify differences using the Levenshtein distance, which attributes equal importance to all amino acid changes in BCR sequences—be it substitutions, insertions, or deletions. While the Levenshtein distance assumes that all mutations influence BCR binding profiles equally, in contrast, the impact of a mutation on PLM embeddings is determined by both the site and nature of the change. In that regard, PLMs offer a new and potentially more expressive approach to analyze immune repertoires during an ongoing immune response.

To examine the extent to which various embeddings capture co-evolutionary relationships among sequences, we leveraged a repertoire of BCR heavy chain sequences originating from 10 GCs from the same lymph node [31]. After processing the sequences to obtain their alignment to germline V and J genes, we assigned them to the same clonal family using standard clonal identification methods [30], i.e. two sequences were grouped together if they had the same V and J genes as well as a CDR3 sequence similarity above 0.9 (Methods Section 4.3). We categorized all sequences based on their V-genes, J-genes, CloneID, and GC of origin (Figure 2A & B). We then assessed the embeddings’ capability to determine whether two BCRs originated from the same or different classes, and we quantified the predictive accuracy in terms of AUC (Figure 2C & D, Table 3). As both the germline gene alignment and the clonal identification rely on variants of the Levenshtein distance, we anticipated that Levenshtein is likely to perform well in these tasks. Not surprisingly, as already hinted by the varying correlations between the studied embeddings and the Levenshtein distance (Table 2), the ability of various PLMs to differentiate B cells from distinct clones or GCs significantly differed across embeddings. While the Levenshtein distance slightly outperformed most embeddings, these differences were only significant when inferring the V-gene – this was expected as the V-gene accounts roughly for the 70% of the sequence, which provides it with an evident advantage for this task. Counter-intuitively, the AbLang model, despite being exclusively trained on antibody sequences, was less proficient in identifying clones and V-genes than the ESM2-based models. However, it excelled in differentiating GCs (Table 3). This once again indicates that each embedding captures distinct information when compared to one another and also when compared to the Levenshtein distance.

**Figure 2:**
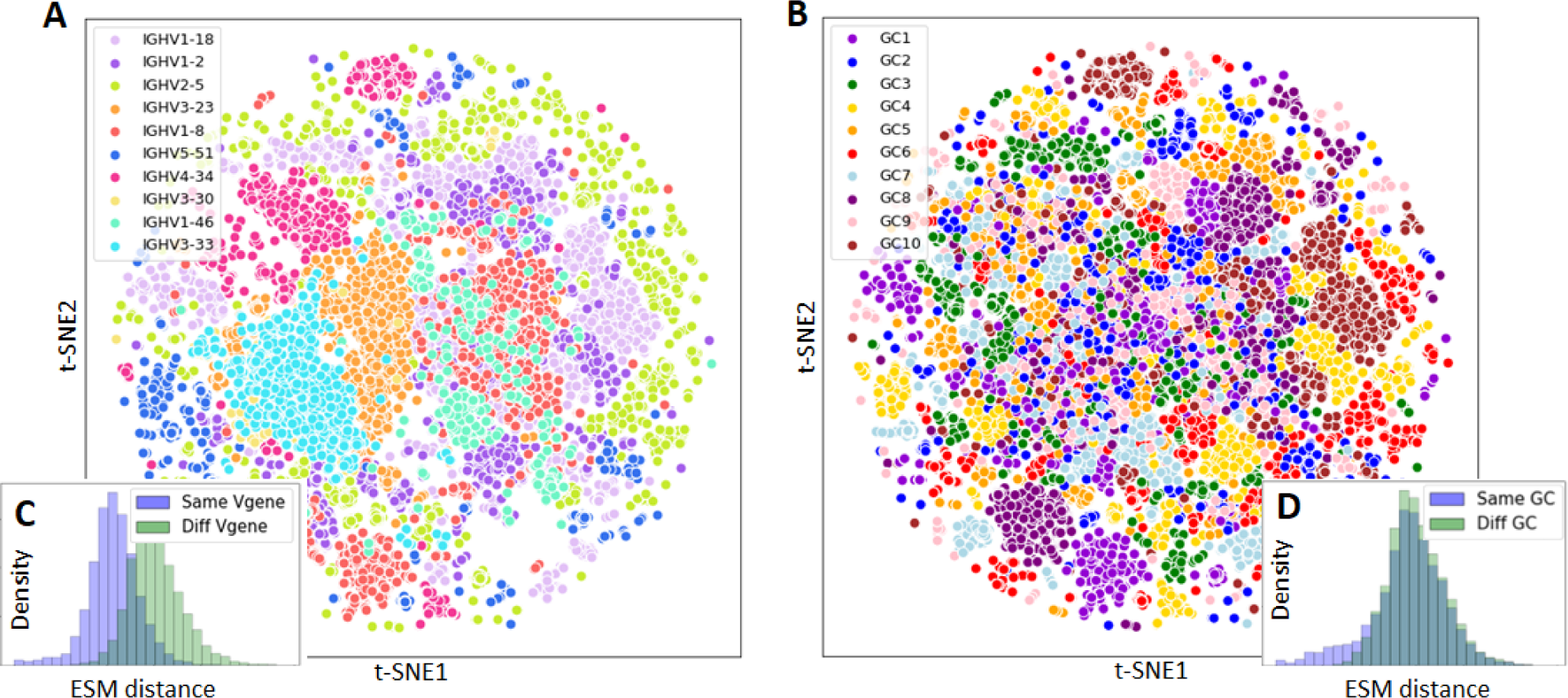
ESM2-650M-L33 t-SNE representation of BCR heavy chains of the GC dataset [31] colored according to (A) their V-gene assignment and (B) their GC of origin. In (C) and (D), we show the distribution of pairwise Euclidean distances in the ESM2 space for BCRs within the same class and across different classes.

**Table 3:** AUC of different paired class differentiation tasks (i.e. do 2 BCRs belong to the same class?) with the Euclidean distance between the embeddings. The classification performance using the Levenshtein distance is also provided for comparison.

|  | V-gene | J-gene | GC | Clone |
| --- | --- | --- | --- | --- |
| Levenshtein | **0.98** | **0.60** | 0.55 | **0.99** |
| AbLang-L12 | 0.84 | 0.59 | **0.57** | 0.96 |
| ESM2-650M-L6 | 0.90 | 0.58 | 0.55 | **0.99** |
| ESM2-650M-L33 | 0.90 | 0.58 | 0.55 | 0.97 |
| ESM2-150M-L30 | 0.89 | 0.56 | 0.54 | 0.97 |
| ESM2-35M-L12 | 0.87 | 0.58 | 0.55 | 0.97 |
| ESM2-8M-L6 | 0.81 | 0.57 | 0.56 | 0.96 |

Next, we investigated and visualized individual B cell clones with Ablang and ESM embeddings. First, we selected a rare clone with a lineage shared among several GCs, typically signaling inter-GC communication events that occur when a B cell migrates from a GC to another [31, 35]. We first constructed its root taking the unmutated V and J germlines and filling the remaining junction region with the consensus sequence of all unique sequences within the considered clone. Then, the phylogenetic evolutionary tree of this clone (Method Section 4.4) revealed clearly distinguishable branches, which roughly correspond to the GC they belong to (Figure 3A). While sequences from GC4 were distinctly separated from the two other GCs, GC10 and GC6 exhibit minor sequence intermixing, which can indicate further B cell recirculation events. An alternative and more plausible explanation might be experimental sequencing errors or sequence mislabeling.

**Figure 3:**
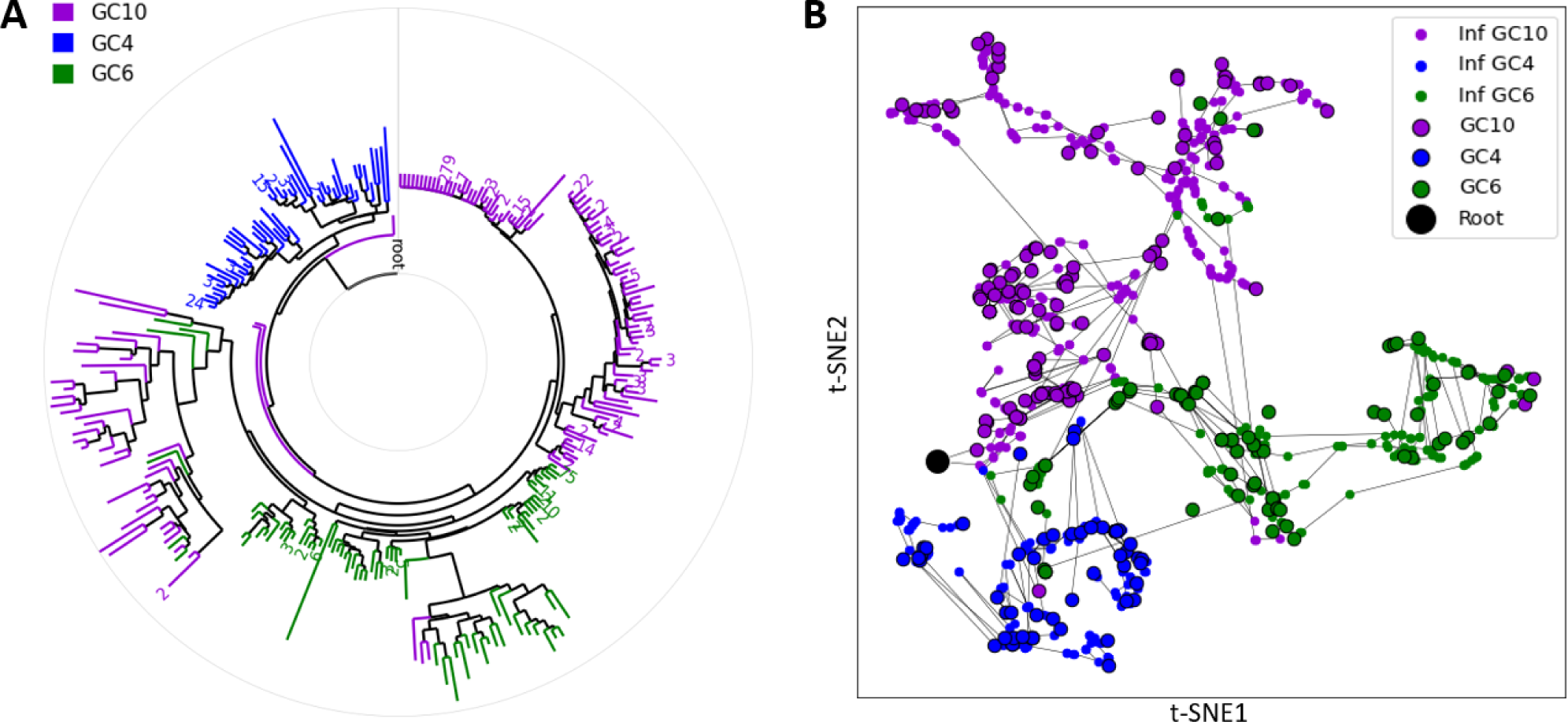
Phylogeny analysis of a B cell clone shared among three germinal centers. (A) Circular phylogenetic tree constructed from a dendrogram of Levenshtein distances between sequences. (B) Visualization of the same B cell phylogeny in the AbLang space. Observed sequences are represented by circled points, while inferred intermediate sequences in the phylogeny are denoted by small points and colored based on their nearest neighbor sequence. Each line represents a single nucleotide mutation between two sequences.

As a comparison, we visualized the same phylogeny using a t-SNE map trained on the BCR Ablang-embedded sequences (Figure 3B, Figure S5), where nucleotide mutations were depicted with a line between neighbors BCR sequences. A simple visualization already revealed meaningful patterns, e.g. B cells from the same GC clustered together and showed correlation with their mutation count from the root, i.e. the number of nucleotide changes that separate a sequence from the root.

Interestingly, a subset of mutations led to larger displacements in the embedding space in terms of Euclidean distance compared to others. While this might be an artifact of t-SNE dimensionality reduction (as seen in Figure S6A, where the same clone is visualized using t-SNE on Levenshtein distances), a systematic analysis of all mutations in the phylogeny from the 50 most abundant clones confirmed that some mutations lead to significantly larger displacements in the embedding space than others. Furthermore, the displacement magnitude induced by a mutation differed significantly depending on whether it was randomly induced or selected during affinity maturation. Here, a random mutation is a random nucleotide change anywhere in the BCR heavy chain, while mutations shaping the inferred phylogenetic tree of expanded clones are referred to as selected. These were inferred during the reconstruction of the B-cell clone phylogeny (Methods Section 4.4).

We observed that AbLang and ESM fundamentally differ in the way they handle random and selected mutations (Figure 4A & B, Figure S7). While AbLang led to a 2-fold increase in the displacement induced by selected mutations compared to random mutations, ESM2 showed the opposite behavior with a 3-fold decrease for the same selected mutations. A similar pattern was found between ESM2-35M, ESM2-150M, and ESM2-650M (Figure 4C). Interestingly, ESM2-650M-L6 did not show a significant difference in terms of the displacement induced by random and selected mutations. Finally, it is worth noting that not all the displacements are equivalent with respect to the Levenshtein distance. For instance, a random mutation has approximately a 20% chance of being silent, resulting in a Levenshtein displacement of zero, whereas other non-silent mutations lead to a displacement of one. As silent mutations are less likely to be selected, this explains the observed slight increase of displacement by selected mutations over random mutations in the Levenshtein space.

**Figure 4:**
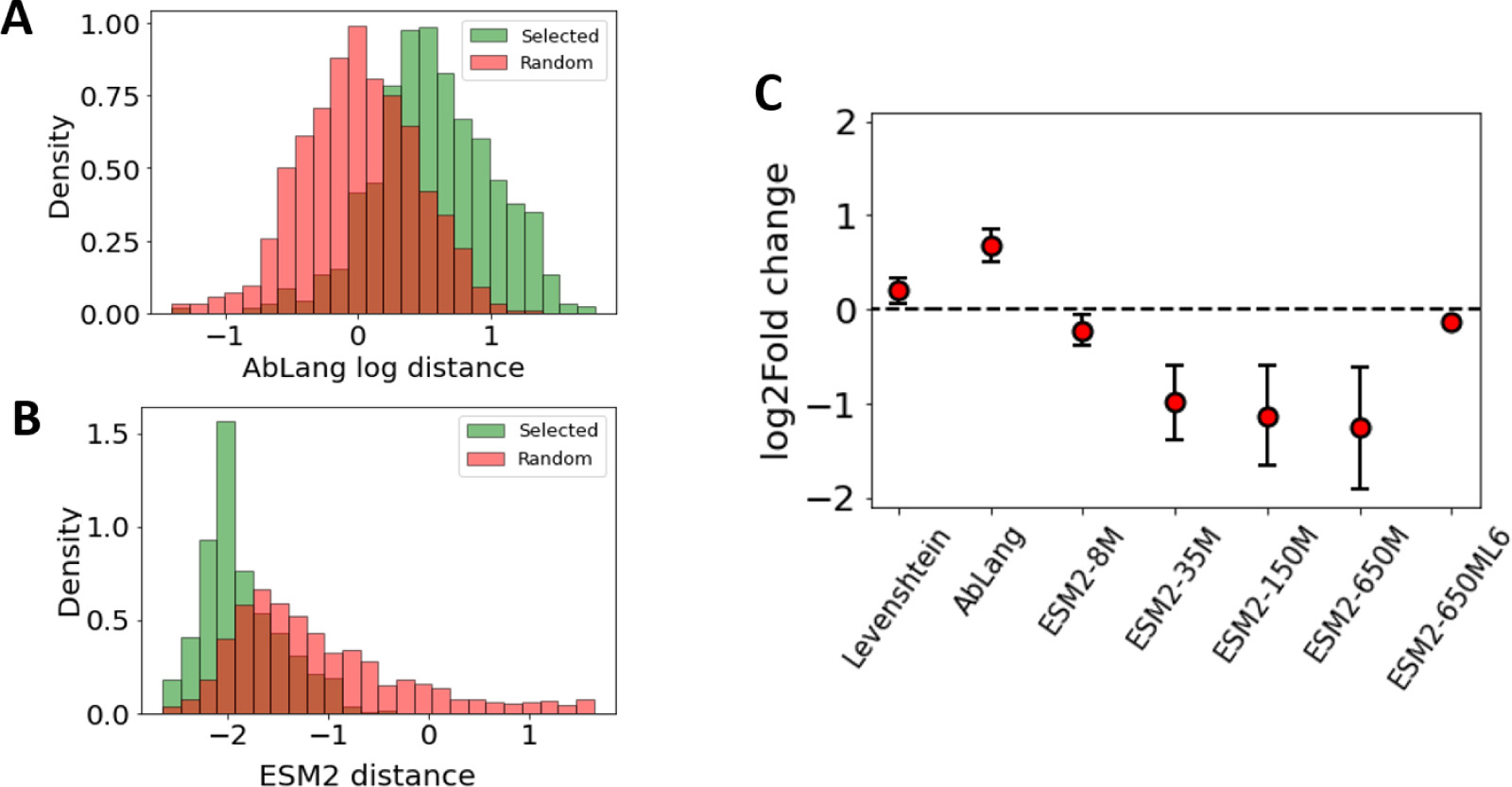
Quantification of displacement in the embedding space after one nucleotide mutation. (A, B) Distribution of Euclidean distance displacements induced by selected and random mutations in the AbLang and ESM2-650M embeddings respectively. (C) log-ratio between the displacements induced by selected and random mutations for different embeddings. The Levenshtein distance is also shown for reference. The values are averaged over the 50 most abundant clones with error bars representing one standard deviation. If no layers are indicated on the PLM *x*-axis, the last layer was used.

While the interpretation of these results is subtle, it is worth noting that the AbLang training dataset comprised a large fraction of naive BCR sequences, i.e., sequences that have not undergone affinity maturation. Thus, we can assume that the embedding space naturally separated naive and mature B cells, as the authors have noted themselves [19]. If this hypothesis is correct, it seems reasonable that selected mutations should induce larger displacement than random ones, as selected mutations are more likely to result in displacements towards the regions of the embedding space where mature sequences lie. On the other hand, since ESM2 has been trained on heterogeneous collections of proteins, the embedding might group sequences by family affiliation rather than by harder-to-predict functional properties, such as affinity. However, further investigation into the properties of the embedded space coupled with data specifically generated to test the properties of interesting regions of this space is needed to better understand the functional implications of displacements in the embedded space.

### 2.3 Comparing generalist and domain-specific embedders for BCR and TCR specificity prediction

It is expected that using machine learning to decode information in adaptive immune receptor repertoires can transform the prediction of immune responses and accelerate the development of vaccines, therapeutics, and diagnostics [36]. Having demonstrated that both Ablang and ESM2 capture important evolutionary information for both BCRs and TCRs, we shifted our focus to examining the predictive accuracy achieved by PLMs in epitope specificity predictions for TCRs and BCRs, both significant open challenges in immunoinformatics [37, 24].

One of the primary limitations of current models for specificity prediction tasks is the scarcity of labeled training data. As a result, techniques leveraging transfer learning approaches [38] and, more recently, extracting embedded representations from extensive models like PLMs are under active exploration [24].

#### TCR predictions

We compared different versions of the ESM2 generalist model and TCR-BERT as featurizers for downstream classification tasks. For this purpose, we borrowed a predictive task from the ImmRep TCR-specificity workshop benchmark, the first public benchmark that evaluated 23 predictive models [37]. The task entails predicting a binary label for binding to the GILGFVFTL epitope given the amino acid sequence of the TCR CDR3-*β* chain. While we focused here on a subset of the ImmRep dataset for a single epitope, we show in the Supplementary Section A.1 that a simple ESM2 embeddings-based classifier with minor modeling effort can compete with the best models from the ImmRep TCR-specificity workshop benchmark [37].

Here, for each embedding-based model, we used a multi-layer perceptron (MLP) with 1, 2, or 3 layers (Methods 4.8) to predict the binary binding label. We report in Table 4 the performance of models selected through hyperparameter optimization, measured with standard metrics for accuracy evaluation: AUC, Accuracy, F1, Precision, and Recall. The architectures and hyperparameters of the selected models are reported in table 7. Interestingly, after optimization of the classifier’s hyperparameters, there was no significant difference between the best ESM2- and TCR-BERT-based classifiers across all metrics considered.

**Table 4:**
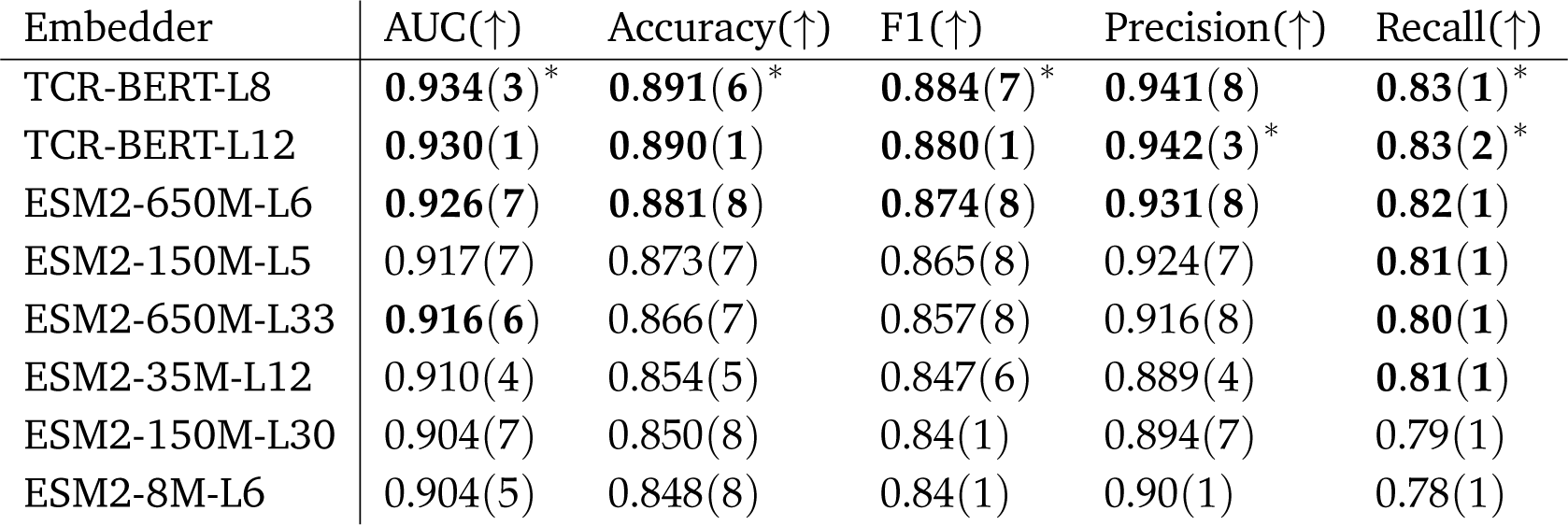
Test metrics evaluated on binary predictions of TCR binding to the GILGFVFTL epitope for hyperparameter-tuned classifiers (selecting on validation AUC) taking different PLM embeddings as input. The highest mean value is indicated with an asterisk, and the standard deviation across Monte Carlo repetitions is indicated in parentheses as the uncertainty on the last significant digit. Values indicated in bold are non-dominated by the best score according to a multiple-testing-corrected Mann-Whitney U-test [39]. Arrows indicate whether higher (*↑*) or lower (*↓*) is better.

| Embedder | AUC(↑) | Accuracy(↑) | F1(↑) | Precision(↑) | Recall(↑) |
| --- | --- | --- | --- | --- | --- |
| TCR-BERT-L8 | **0.934(3)\*** | **0.891(6)\*** | **0.884(7)\*** | **0.941(8)** | **0.83(1)\*** |
| TCR-BERT-L12 | **0.930(1)** | **0.890(1)** | **0.880(1)** | **0.942(3)\*** | **0.83(2)\*** |
| ESM2-650M-L6 | **0.926(7)** | **0.881(8)** | **0.874(8)** | **0.931(8)** | **0.82(1)** |
| ESM2-150M-L5 | 0.917(7) | 0.873(7) | 0.865(8) | 0.924(7) | **0.81(1)** |
| ESM2-650M-L33 | **0.916(6)** | 0.866(7) | 0.857(8) | 0.916(8) | **0.80(1)** |
| ESM2-35M-L12 | 0.910(4) | 0.854(5) | 0.847(6) | 0.889(4) | **0.81(1)** |
| ESM2-150M-L30 | 0.904(7) | 0.850(8) | 0.84(1) | 0.894(7) | 0.79(1) |
| ESM2-8M-L6 | 0.904(5) | 0.848(8) | 0.84(1) | 0.90(1) | 0.78(1) |

**Table 5:** Test metrics evaluated on predictions of antibody binding to a SARS-CoV-2 epitope for hyperparameter-tuned regression models (selecting on validation R^2^) taking different PLM embeddings as input. Values indicated in bold are non-dominated by the best score according to a multiple-testing-corrected WMU test. Arrows indicate whether higher (*↑*) or lower (*↓*) is better.

| Embedder | MSE (↓) | MAE (↓) | R<sup>2</sup> (↑) |
| --- | --- | --- | --- |
| ESM2-650M-L6 | **0.119(1)** | **0.2659(9)** | **0.455(7)** |
| ESM2-650M-L33 | 0.123(1) | 0.271(2) | 0.438(6) |
| AbLang-L12 | 0.127(1) | 0.276(3) | 0.419(4) |
| ESM2-35M-L12 | 0.129(1) | 0.279(2) | 0.413(5) |
| ESM2-150M-L30 | 0.1305(9) | 0.283(2) | 0.404(4) |

**Table 6:** Optimal hyperparameter choices for BCR affinity regression models (Multilayer perceptron) taking embeddings from each PLM tested. The layers correspond to the output dimension of each hidden layer and the number of parameters (#Param.) takes into account each linear layer matrix and bias term.

| PLM | Best Hyperparameter Choice for Multilayer Perceptron |  |  |  |  |
| --- | --- | --- | --- | --- | --- |
|  | PLM Dim. | Layers | #Param. | Learning Rate | Dropout Rate |
| ESM2-650M-L6 | 2560 | [128, 32] | 331,936 | $1 \times 10^{-4}$ | 0.0 |
| ESM2-650M-L33 | 2560 | [512, 128] | 1,376,896 | $1 \times 10^{-4}$ | 0.0 |
| ESM2-35M-L12 | 960 | [512, 128] | 557,696 | $1 \times 10^{-4}$ | 0.0 |
| ESM2-150M-L30 | 1280 | [512, 128] | 721,536 | $1 \times 10^{-3}$ | 0.0 |
| AbLang-L12 | 1536 | [512, 128] | 852,608 | $1 \times 10^{-4}$ | 0.05 |

**Table 7:**
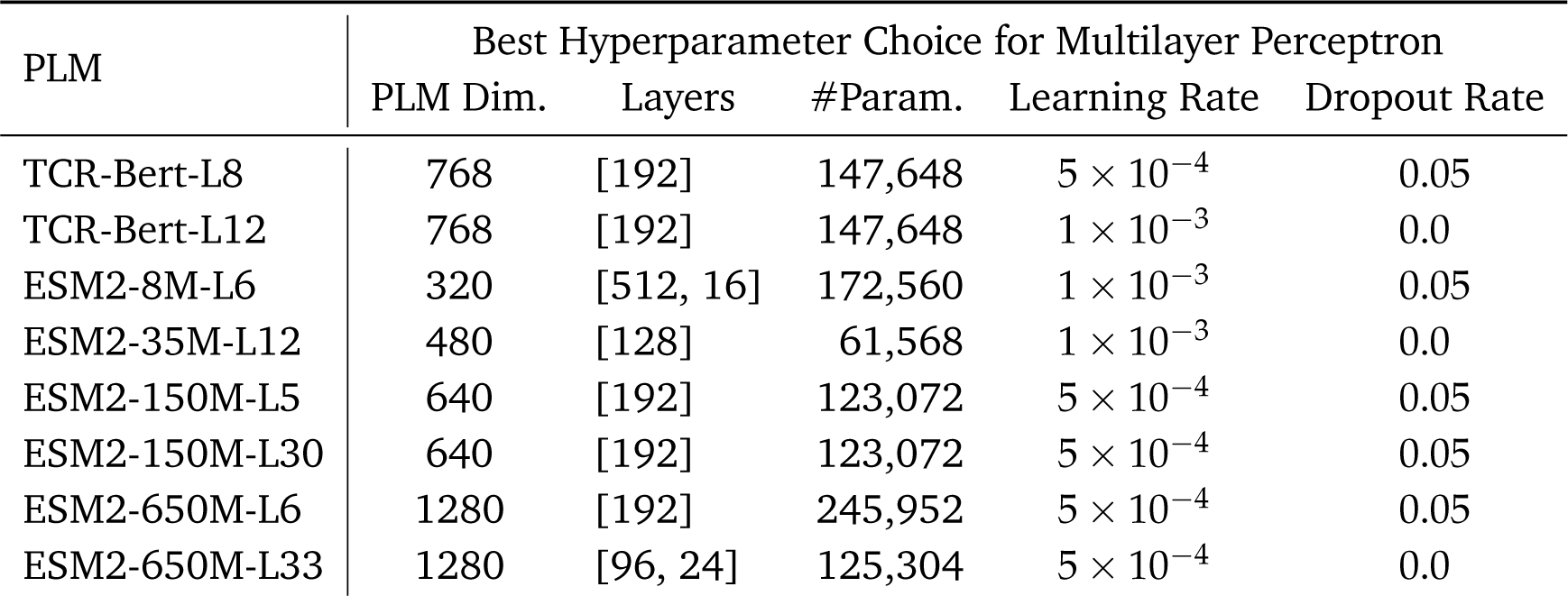
Optimal hyperparameter choices for TCR binding classifiers (Multilayer perceptron) taking embeddings from each PLM tested. The layers correspond to the output dimension of each hidden layer and the number of parameters (#Param.) takes into account each linear layer matrix and bias term.

| PLM | Best Hyperparameter Choice for Multilayer Perceptron |  |  |  |  |
| --- | --- | --- | --- | --- | --- |
|  | PLM Dim. | Layers | #Param. | Learning Rate | Dropout Rate |
| TCR-Bert-L8 | 768 | [192] | 147,648 | $5 \times 10^{-4}$ | 0.05 |
| TCR-Bert-L12 | 768 | [192] | 147,648 | $1 \times 10^{-3}$ | 0.0 |
| ESM2-8M-L6 | 320 | [512, 16] | 172,560 | $1 \times 10^{-3}$ | 0.05 |
| ESM2-35M-L12 | 480 | [128] | 61,568 | $1 \times 10^{-3}$ | 0.0 |
| ESM2-150M-L5 | 640 | [192] | 123,072 | $5 \times 10^{-4}$ | 0.05 |
| ESM2-150M-L30 | 640 | [192] | 123,072 | $5 \times 10^{-4}$ | 0.05 |
| ESM2-650M-L6 | 1280 | [192] | 245,952 | $5 \times 10^{-4}$ | 0.05 |
| ESM2-650M-L33 | 1280 | [96, 24] | 125,304 | $5 \times 10^{-4}$ | 0.0 |

As previously discussed [22], using early layers can lead to better performance in downstream tasks. For ESM2-based models, we observed that training on embeddings from an early layer of the largest model we considered (ESM2-650M-L6), yields significantly better performance for several metrics compared to classifiers trained on embeddings from the last layer (ESM-650M-L33). For TCR-BERT, however, we observed no significant difference between the last layer of TCR-BERT (L12) and the layer recommended for downstream affinity prediction (L8) [22], although the latter has a much lower AUC standard deviation which is a desirable feature in terms of performance guarantees. One final feature of note in Table 4 is the lack of significant differences in recall among models based on six of the eight embeddings we considered, in stark contrast with the other metrics. This is explained by the empirical observation that a larger fraction of the positively labeled (binding) samples is close to the models’ decision boundaries, making them more sensitive to statistical noise. A potential reason for this observation is the design of the training set where the positive samples are those with certain properties that lead to the high binding affinity, while the negative samples are unrelated randomly chosen TCRs. This makes it less likely for individual negative samples to decisively define the decision boundary.

Finally, a possible factor affecting the embedding’s performance could be the size of the MLP used to classify TCR binders, with larger models potentially able to capture more relevant features for downstream tasks. Interestingly, in Supplementary Section A.2, we show that there was no visible dependence of optimized performance on parameter count for TCR binding predictions.

#### BCR predictions

We turned next to BCR binding prediction tasks, which share similar biochemical underpinnings as TCR specificity prediction, but also present important differences from a machine learning point of view. We considered the AlphaSeq dataset [40], which provides continuous measurements of antibody affinity to an epitope of the SARS-CoV-2 spike protein. We defined the predicting affinity task as a continuous regression task, rather than a binary classification as we did for TCR specificity prediction. Another important difference is in the size of the sample, which comprises 154,029 samples including replicates, and 71,415 paired heavy and light chains. This dataset is thus approximately 37 times larger than the TCR dataset.

Similarly, as for the TCR binding prediction, we used ESM2 as the state-of-the-art general protein PLM, and compared its performance with AbLang, the best-in-class antibody-specific model [24]. For each BCR, we embedded the paired heavy and light chains separately. Then, we concatenated the embeddings and processed them with a shallow MLP (2 to 4 layers and 500K - 1.5M parameters, Methods Section 6) to obtain an affinity prediction. We optimized the regression models separately and selected the optimal hyperparameter set based on a held-out validation set.

We present the performance of the optimized classifiers in Table 5, whose architecture and hyperparameters are described in Table 6. Here, as for TCRs, the best ESM2 embedding was obtained from an early layer (layer 6) of the largest embedding (650M parameters). Regression models constructed using these embeddings significantly outperformed both the other ESM2 embeddings and the AbLang-based embeddings. The second-best-performing model on all metrics is ESM2-650M-L33, which is the same model as the best-performing model, but with embeddings extracted from the final layer. This second model also significantly outperformed AbLang in terms of MSE (Mean Squared Error) and R^2^ (coefficient of determination), but did not statistically dominate it in terms of MAE (Mean Absolute Error). As we observed with TCRs, the size of the MLP did not affect the performance of the BCR binding predictions (Supplementary Section A.2).

In contrast with the analogous TCR binding prediction models where the best ESM2 models had comparable performance with domain-specific TCR-BERT, Ablang was clearly dominated by ESM2 here. A possible straightforward explanation could be that AbLang encodes less relevant information about antibody binding compared to the best ESM2 embedding, while for TCR-related tasks, a similar amount of information is captured by ESM2 and TCR-BERT embeddings. However, an alternative explanation might be that ESM2 encodes richer protein information compared to the domain-specific models. Yet, the limited sample size of the TCR classification task, which is 37 times smaller than AlphaSeq, might prevent the classifier from effectively differentiating relevant from non-task-specific features, limiting ESM2 accuracy.

To test this hypothesis, we considered multiple randomly down-sampled versions of the AlphaSeq dataset. For each dataset, we trained a regression model by selecting hyperparameters optimized specifically for each dataset size, for both AbLang-L12 and the superior ESM2 model, ESM2-650M-L6. As expected and shown in Fig. 5, the performance of both PLM embedding-based regressors deteriorated with decreasing training data sizes. The figure shows that there is a clear gap between the two embeddings in the high-data regime, where ESM2-650M-L6 outperformed AbLang. Furthermore, ESM2-650M-L6 performance has an earlier inflection point while AbLang performance decreased more steadily with scarcer data, as a result of which, both models have comparable performances in the low-data regime. This observation supports the hypothesis that both PLMs encode similarly predictive *learnable* information when training data is limited. However, when more training data is available, the larger ESM2-650M-L6 encodes more relevant information than AbLang, at least, for the task we have inspected. This increase in disparity in the rich-data size regime could be attributed to the differences in their embeddings’ dimensions (1280 compared to 768 for Ablang, Table 1). ESM2, having higher dimensionality, may require more training data to reach its maximum disparity.

**Figure 5:**
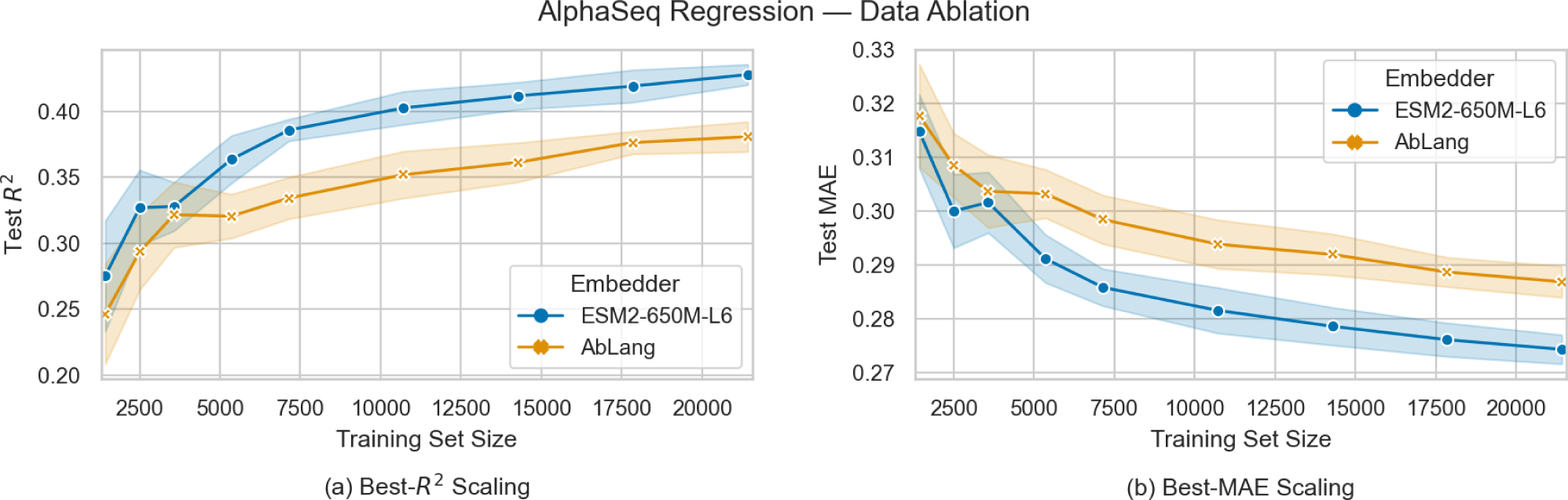
Test performance of hyperparameter-tuned regression models taking AbLang-L12 and ESM2-650M-L6 embeddings as input as a function of the training set size. (a) Test R^2^ scores of models with the best validation R^2^. (b) Test MAE scores of models with the best validation MAE.

## 3 Discussion

The profound impact and utility of Protein Language Models (PLMs) in bioinformatics [10] and immune-related applications [24] is beyond dispute. However, as new PLMs continuously emerge, it remains unclear which factors are essential to consider when utilizing them. Our study has shown that the model’s size, the magnitude of the training dataset, and the selection of layers, all play crucial roles in the model’s ability to capture relevant biological features in immunological tasks. For instance, ESM2-650M embeddings, with their higher dimensionality (1280 for all its layers compared to 768 for Ablang, Table 1), outperforms domain-specific PLMs only when a sufficient amount of training data is available. Concerning the model’s size, ESM2 consistently shows enhanced performance as the parameter size grows, aligning with the results seen in other related tasks where it was evaluated [12, 15]. We also found that the last layer of PLMs is not necessarily the one yielding the highest performance, and thus, the layer number should be considered as a hyperparameter that needs to be optimized for each downstream task.

Regarding BCRs, their unique evolutionary processes to maximize the binding affinity to varying targets set them apart from most proteins [41]. Indeed, the close relationship between an antibody and the germline sequence of the naive B cell from which it has evolved, along with the subsequent random mutations it has undergone, collectively determine its specificity and binding profile. The distinct selective pressures experienced within germinal centers further shape BCR repertoires in a manner that presents challenges for generalist PLMs to accurately represent. For example, we observed that selected mutations had a greater impact on AbLang embeddings compared to random mutations, presumably because they enhanced the binding profile toward a specific target. However, this pattern was reversed for ESM2 embeddings (Figure 4). While a comprehensive exploration of the meaning of displacements in the embedding space requires specialized techniques from interpretable deep learning and falls beyond the scope of this work, this different behavior suggests that AbLang and ESM2 organize BCR sequences in the embedding space in significantly distinct ways.

However, understanding how this different organization impacts the predictive performance of different embeddings on various downstream tasks is not trivial. While it might be naively expected that domain-specific models should outperform generalist models on tasks with high evolutionary specificity, it was demonstrated that ESM1 [12], the predecessor of ESM2, performed comparably to AbLang on various antibody-related tasks, including paratope and antigen binding prediction [24]. Adding to the comparison, we have shown here that ESM2 clearly outperforms AbLang on antigen-binding tasks. On the other hand, ESM1 underperformed AbLang on tasks requiring high specificity, such as antibody discovery and cell state prediction. Therefore, AbLang may still be a competitive choice compared to ESM2 for these tasks.

An intriguing question that our study does not address is to better understand which information gets selected in the low-data regime. Even without a thorough interpretation of the information captured by each embedding, assessing the generalization of their predictions on unseen data and the stability of predictions across subsampled training datasets could yield valuable insights into the information utilized by the downstream regressors. Of particular interest is whether there are specific ESM2 embedding features that are only reliably leveraged in the high data regime, leading to its improved performance, or whether the low-data regressors consistently make global but shallow use of the available ESM2 embedding features, suggesting a more robust and generalizable approach in scenarios with limited data.

Finally, in the context of modeling the evolution of antibodies during an immune response, the utilization of PLMs represents a promising approach for predicting the impact of mutations [42]. The limited accuracy of current antibody-antigen binding prediction models poses substantial challenges in the field of antibody design [7]. Rather than explicitly simulating the molecular binding between the antibody with its antigen, PLMs offer an alternative approach to train binding prediction models. Current modeling efforts to simulate an immune response to a known antigen cannot evaluate the influence of point mutations on BCR sequences [43, 44]. This limitation hinders their applicability in vaccine design or infectious disease research, where faithful simulations of repertoire adaptation to external threats are needed. In this context, PLMs could facilitate more realistic germinal center simulations [43, 45, 46, 47] by predicting which amino acids are more likely to undergo mutations.

Similar potential is expected in the context of T cells, where the development of T cell-based immunotherapies is currently hampered by our limited ability to predict cross-reactive binding events, i.e. events where natural or engineered T cells bind to off-target sites, potentially triggering inflammatory responses at healthy sites. Although current PLMs cannot predict the binding affinity to unseen epitopes and antigens, their predictive accuracy for numerous tasks is advancing rapidly. Aid by emerging high-throughput single-cell technologies that may enable the swift measurement of BCRs and TCRs binding affinity to extensive collections of epitopes and antigens, there is promising potential to revolutionize the clinical application of both B and T cell-based therapies.

## 4 Methods

### 4.1 Datasets

#### TCR specificity

The TCR dataset was provided by the organizers of the ImmRep 2022 TCR-epitope specificity workshop [37], and was downloaded from https://github.com/viragbioinfo/ImmRep_2022_TCRSpecificity. We restrict our focus to TCRs with binding information for the GILGFVFTL epitope. This selection deviates from the broader ImmRep goal to test various epitopes, and is attributed to the fact that TCR-BERT only embeds TCR sequences, not epitopes. A mix of positive and negative TCRs for each epitope was provided in the test data. By concentrating on a single epitope, our sample size is reduced to 4067 sequences, of which 680 are positively labeled. While the dataset contains paired information on the *α* and *β* chains, we retained only the CDR3-*β* chain to construct embeddings due to TCR-BERT limitations.

#### Antibody specificity

The antibody covid specificity Dataset (AlphaSeq) [40] was downloaded from https://zenodo.org/record/5095284. AlphaSeq consists of 71,415 paired heavy and light chains, with continuous measurements of their affinity to an epitope of the SARS-CoV-2 spike protein. The sequences were generated by introducing artificial mutations (up to three) in the CDRs of four known binders of the SARS-CoV-2 spike protein.

#### Germinal center BCRs

The collection of B cell sequences originating from 10 individual germinal centers from a single human lymph node [31] was downloaded from https://vdjserver.org/commun ity/8899006209436478995-242ac118-0001-012. Germline V and J genes assignments were already included in the datasets, and the clone phylogenies were inferred following the approach described in Method 4.3. Sequences with NaN values in V or J gene segments were removed from the study. Sequences are between 300 and 340 nucleotides long, and the V and J segments comprise roughly 70% and 15% of the sequences, respectively.

### 4.2 Levenshtein distance

The Levenshtein distance [29], defined as the minimum number of edits required to transform one sequence into another, is a common metric to quantify sequence similarity. To reduce the bias caused by differences in sequence length, we use the normalized Levenshtein distance [48] that incorporates the length of both sequences and satisfies the triangle inequality. Given two strings, *s*_1_ and *s*_2_, and the Levenshtein distance between them, *Lev*(*s*_1_, *s*_2_), the normalized Levenshtein distance *Lev*_norm_ is defined as:

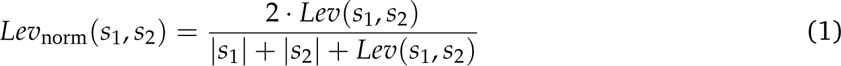

### 4.3 Identifying germinal center B-cell clones

Following on the recommendations of [30], we first grouped B cells together only if they had the same V and J genes. For each obtained group, we computed the pairwise normalized Levenshtein distance between each junction sequence in that group, and applied the Hierarchical Agglomerative Clustering (HAC) algorithm [49, 50] to cluster the BCRs into different clonal groups. We used the complete-linkage clustering criterion, which begins by putting each sequence on its own cluster, and then sequentially combines smaller clusters into larger ones until all elements are in one single cluster. This method generates a dendrogram, illustrating the sequence of cluster mergers and the distance at which each merger occurred. By setting an appropriate threshold, we can define individual clusters as all the clusters that have not been merged up to that distance. In this study, we chose as threshold the distance to the nearest distribution of negation sequences with a tolerance of 0.03% [30], corresponding to a threshold distance of 0.1. Here, negation sequences refer to BCR sequences with unrelated phylogenies, typically sampled from unrelated individuals.

### 4.4 Inferring the mutation phylogeny of B-cell clones

We used mutation phylogenetic trees to visualize and reconstruct the evolution of the BCR sequences in a given clone. In such representation, each founder cell defines the unmutated germline of a new tree, newly acquired mutations are represented as downstream nodes (Figure S4A), and leaves represent observed sequences [43].

To construct the tree, we first defined the root by taking the unmutated V and J germlines and filling the remaining junction region with the consensus sequence of all unique sequences within the considered clone. Then, the grouped sequences from each clone were aligned with ClustalW [51]. The alignment was necessary to define distances between each sequence, from which we inferred a tree with a hierarchical clustering approach. The most similar sequences were grouped together and progressively aggregated with other groups until the root node (maximum distance) was reached (Neighbour Joining method in ClustalW). Then, we associated a sequence to each bifurcation in the dendrogram, defined as the consensus sequence of all the leaves below it. Finally, the final tree was constructed by linking bifurcations together and by introducing mutations between them, or by merging them when having the same sequence. These inferred sequences, or intermediate nodes in the tree, represent the BCR of parent cells that existed but later underwent additional mutations and are not present in the repertoire at the sequencing snapshot. The compilation of mutations linking these inferred sequences together defines a set of *selected* mutations that we can leverage to evaluate the sensitivity of PLMs to these mutations.

This method to infer sequences was preferred over Bayesian Maximum likelihood estimation approaches [52, 53] or Markov chain Monte Carlo sampling [54], as these heavily rely on the infinite sites assumptions, according to which every mutation occurs at a previously not mutated site. While this assumption is reasonable at a genome-wide scale, in the context of shorter junctions and somatic hypermutation rates, it is often not verified, as we have found in our data (Figure S4B).

### 4.5 Visualizing phylogenetic trees

Circular visualization of Hierarchical dendogram were implemented with pyCirclize [55] (https://github.com/moshi4/pyCirclize). Mutation phylogenetic trees were plotted with Graphviz (https://pypi.org/project/graphviz/).

### 4.6 Implementation details of PLMs

All embeddings are extracted using PyTorch [56] implementations of the aforementioned PLMs. For each model, we tokenize the protein sequences using the tokenizers provided by the authors and obtain token-wise vector representations using a forward pass of their encoders. These embedded vector representations of variable-length amino acid sequences are aggregated to produce fixed-size representations using mean pooling, which are then used as global sequence representations. Model-specific details follow.

#### ESM2

We use the PyTorch-Hub implementation of ESM2 models [15] (https://github.com/faceb ookresearch/esm) and rely on the readily-available repr_layers parameter of the esm.ESM2.forward method to collect the token embeddings, i.e. the amino acid-based embeddings, at any given internal layer of the model. Mean pooling is achieved by truncating each padded vector sequence at its correct length before computing the per-sequence mean.

#### TCR-BERT

We use the HuggingFace [57] implementation of the base version of TCR-BERT (trained exclusively using amino acid masked language modeling), which is available under the label wukevin/tcr-bert-mlm-only on the HuggingFace Hub. Another version of the model, fine-tuned on a TCR binding prediction task, is available but raises the risk of having part of our test set exposed as its training data. We obtain mean-pooled embeddings with the SentenceTransformers library [58], which handles mean pooling automatically. Early layers were obtained by editing the HuggingFace model internals (truncating the BertModel.encoder.layer implementing the sequence of 12 BertLayer) before being passed to the sentence_transformers.SentenceTransformer object.

#### AbLang

The AbLang [19] authors have made their model available as a standalone Python package(ht tps://github.com/oxpig/AbLang) which readily provides embedding computation through its main interface. We use this interface directly (calling ablang.pretrained objects with mode=’seqcoding’). AbLang comprises two models, optimized for modeling antibody heavy and light chains, respectively. We use each model to obtain a mean-pooled encoding of each BCR separately, which we then concatenate to obtain a global antibody representation.

### 4.7 Model architecture for antibody specificity prediction

We evaluate PLM-based antibody affinity predictions through the lens of the AlphaSeq dataset [40], which provides a continuous label for the estimated binding affinity of antibodies to a specific SARS-CoV-2 epitope, including several replicate measurements for many sequences. We use PLMs to compute fixed-size embeddings of the antibody protein sequences, which reduces the task of predicting the affinity to a supervised regression problem, mapping embedding vectors to a continuous number. Given that AbLang provides separate embeddings for the antibody heavy and light chains, we obtain antibody-level representation by concatenating the two chain vector representations. To ensure a fair comparison with ESM2-based embeddings, we replicate this process for ESM2 models. Specifically, we individually embed each chain and then concatenate them both into a single vector.

We use multi-layer perceptrons of 2, 3, or 4 layers to map the high-dimensional sequence representations to a single continuous label. We use dropout regularization after the internal layers and before the non-linear ReLU activation functions, but neither dropout nor an activation function is applied to the output of the final layer. The model parameters are trained by optimizing the mean-squared error (MSE) loss using the Adam optimizer [59] until a minimum validation loss is achieved, based on an early-stopping algorithm with a patience of 5 steps. Given the large size of the dataset, we set the batch size to a fixed value of 128 to accelerate training and do not consider lower values.

Model hyperparameters are tuned using a random search with Monte-Carlo cross-validation with six repetitions. For each Monte Carlo replicate, 20% of the data is held out for testing and 20% of the remaining data is held out for validation. Hence, the data is split as follows, 64% for training, 20% for testing, and 16% for validation. For each training, model weights are randomly initialized and we use the training data to estimate the coordinate-wise mean and standard deviation of the embedding distribution and standardize training, validation, and test data with these estimates.

In each model, we specify the architecture by specifying the dimensions of the internal layers, while the input dimension is determined by the dimensionality of the embedding vectors used (see Table 1), multiplied by two due to the concatenation, and the output dimension is always 1. The explicit range of random parameters is listed below.

- Learning Rate: [10*^−^*^4^, 10*^−^*^3^, 10*^−^*^2^].
- Dropout Rate: [0, 0.05, 0.1, 0.3].
- Batch size: fixed to 128.
- Hidden Layers: [(128, 32), (512, 128), (1024, 512, 512), (512, 512, 512, 512)].

In Figure 6, we show the ten best hyperparameter (HP) configurations of each test PLMs with a box-plot generated with six samplings [60], ranked by their mean *R*^2^ test score. Strikingly, the best AbLang hyperparameter configuration comes at the 11th position, reinforcing our confidence in the superiority of ESM2 embeddings.

**Figure 6:**
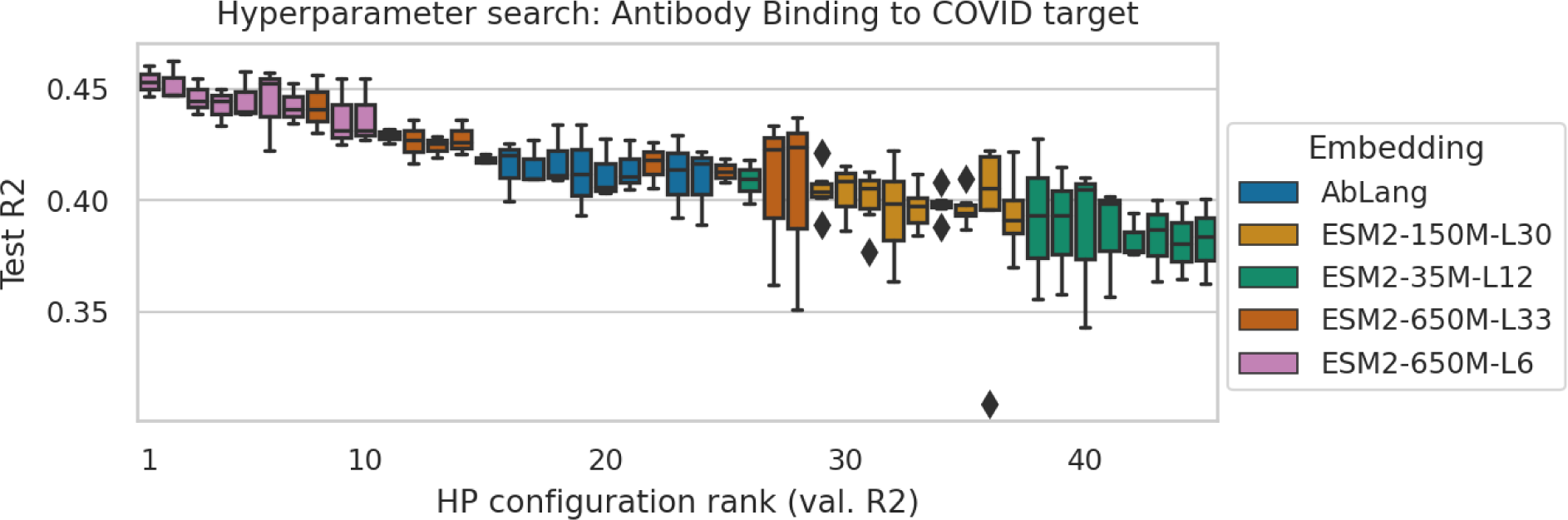
Test *R*^2^ score distributions of the 10 best TCR binding classifier hyperparameter (HP) configurations for each PLM we tested. We rank hyperparameter choices based on their mean validation *R*^2^ score across six Monte Carlo replicates, which we visualize with a Box-plot [60]. Each position on the horizontal axis corresponds to a choice of PLM and HP.

### 4.8 Model architecture for TCR binding prediction

The ImmRep benchmark [37] sets up TCR-epitope binding prediction as a binary classification task where a TCR-epitope pair is labeled as positive if the TCR recognizes the epitope, while negatively labeled pairs indicate non-recognition. In the TCR task, we consider in this manuscript, we identify the individual epitope with the most positively labeled samples and simplify the task to a binary TCR classification problem. We take the CDR3-*β* segment of the TCR and use protein-PLMs as sequence featurizers, mapping variable-length amino-acid sequences to fixed-size vectors. The task then reduces to a standard supervised binary classification problem, mapping a vector of features to a binary label. Given the lack of obvious internal structure or invariances in the embedding spaces we consider, we use multi-layer perceptrons to map the high-dimensional sequence representations to a single binary label, normalized with a sigmoid function. We use dropout regularization after internal layers and before non-linear ReLU activation functions. The model parameters are trained by optimizing the binary cross-entropy loss using the Adam optimizer [59]. Training continues until a minimum validation loss is achieved. We assume that a minimum has been reached if there is no observed improvement after training for 20 epochs.

Model hyperparameters are tuned using a random search with Monte-Carlo cross-validation with six repetitions. For each Monte Carlo replicate, 20% of the data is held out for testing and 20% of the remaining data is held out for validation. Hence, the data is split as follows, 64% for training, 20% for testing, and 16% for validation. For each training fold, model weights are randomly initialized and we use the training data to estimate the coordinate-wise mean and standard deviation of the embedding distribution and standardize training, validation, and test data with these estimates.

In each model, the input dimension is determined by the dimensionality of the embedding vectors used and the output dimension is always 1. We specify the architecture by specifying the dimensions of the internal layers which we tune by defining a scaling factor *λ*. The explicit range of random hyperparameters is listed below.

- Learning Rate: [10*^−^*^4^, 5 *×* 10*^−^*^4^, 10*^−^*^3^, 5 *×* 10*^−^*^3^, 10*^−^*^2^].
- Dropout Rate: [0, 0.05, 0.1, 0.15, 0.2, 0.3].
- Batch Size: [8, 16, 32, 64, 128].
- Hidden Layers: [(*λ ×* 64), (*λ ×* 32, *λ ×* 8), (*λ ×* 64, *λ ×* 8), (*λ ×* 256, *λ ×* 8), (*λ ×* 32, *λ ×* 32, *λ ×* 8), (*λ ×* 64, *λ ×* 32, *λ ×* 8)].

For each value of the hidden-layer-scale factor *λ* = 1, 2, 3, we sample 32 random choices of hyperparameters on which we run our six Monte-Carlo cross-validation replicates. The best models used to report scores are selected based on the mean AUC validation score across Monte Carlo replicates. We report test AUC score distributions among the 10 best hyperparameter choices in Figure 7. The actual top-performing models’ hyperparameter choices are specified in Table 7.

**Figure 7:**
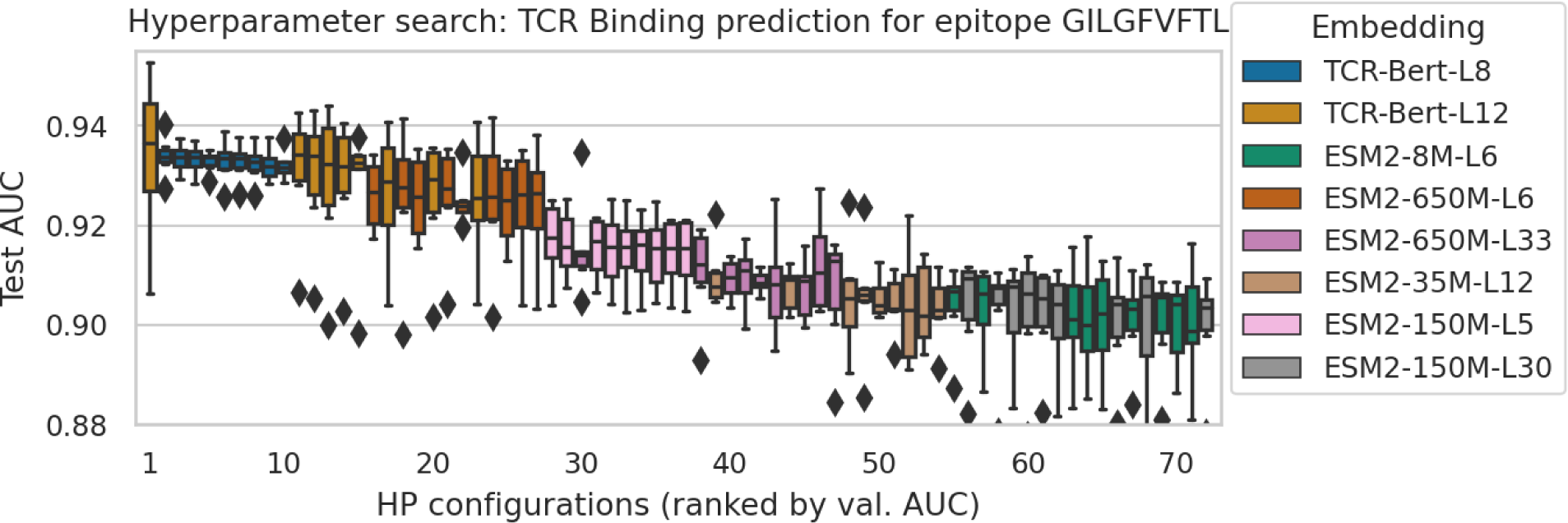
Test AUC score distributions of the 10 best TCR binding classifier hyperparameter (HP) configurations for each embedding we tested. We rank hyperparameter choices based on their mean validation AUC score across six Monte Carlo replicates, which we visualize with a Box-plot [60]

Note that the range of parameters scanned in this experiment is more comprehensive than for antibody affinity regression. This is the result of our observation of the lack of significant difference between the best TCR-BERT- and ESM2-based models, which led us to perform a second random search with expanded parameter ranges and to introduce the layer-width scaling factor *λ*.

## Competing Interests

There is NO Competing Interest.

## Author Contributions Statement

A.P. and N.D. performed computations and wrote the manuscript, under the supervision of M.R.M. A.W. and S.G. performed computation of supplementary section A, under the supervision of M.R.M. and J.B. All authors have read, reviewed, and agreed to the published version of the manuscript.

## Acknowledgments And Funding

This work was supported by the Swiss National Science Foundation Grant No 192128, and by the European Union’s Horizon 2020 research and innovation program under grant agreements No 765158 (COSMIC), No 813545 (HELICAL), and No 826121 (iPC).

## Supplementary Materials

## A Supplementary Text

### A.1 Comparison of the optimal ESM2-based models for paired ***α*** and ***β*** TCR chain sequences

Here, we compare the performance of hyperparameter-optimized ESM-based models to the ImmRep benchmark submissions. As this benchmark used single-cell paired *α* and *β* chains, in this version, we embedded the full TCR sequence (both *α* and *β* chains) and concatenated them together with the embedding of the epitope amino-acid sequence, creating a large vector which is processed by an MLP to predict a binding probability. The best model was found to be a 2-hidden-layer MLP with 512 neurons on each layer, trained with the Adam optimizer [59] with a learning rate of 10*^−^*^4^ and a dropout rate of 0.1. Figure S1 shows that we obtain competitive performance in this task with minor modeling effort.

We also evaluate ESM on an epitope ranking task, a second task proposed in the ImmRep 2022 benchmark. Namely, for each TCR, 17 epitopes were ranked based on model predictions from most to least likely to bind that TCR, and the rank of the ground truth is averaged across all TCRs for a given epitope. With a ROC AUC of 0.83 for the TCR classification task and an average rank of 3.8 for the epitope ranking task, our ESM2 embeddings-based classifier ranks among the top-performing models on the ImmRep benchmark (Figure S1).

**Figure S1:**
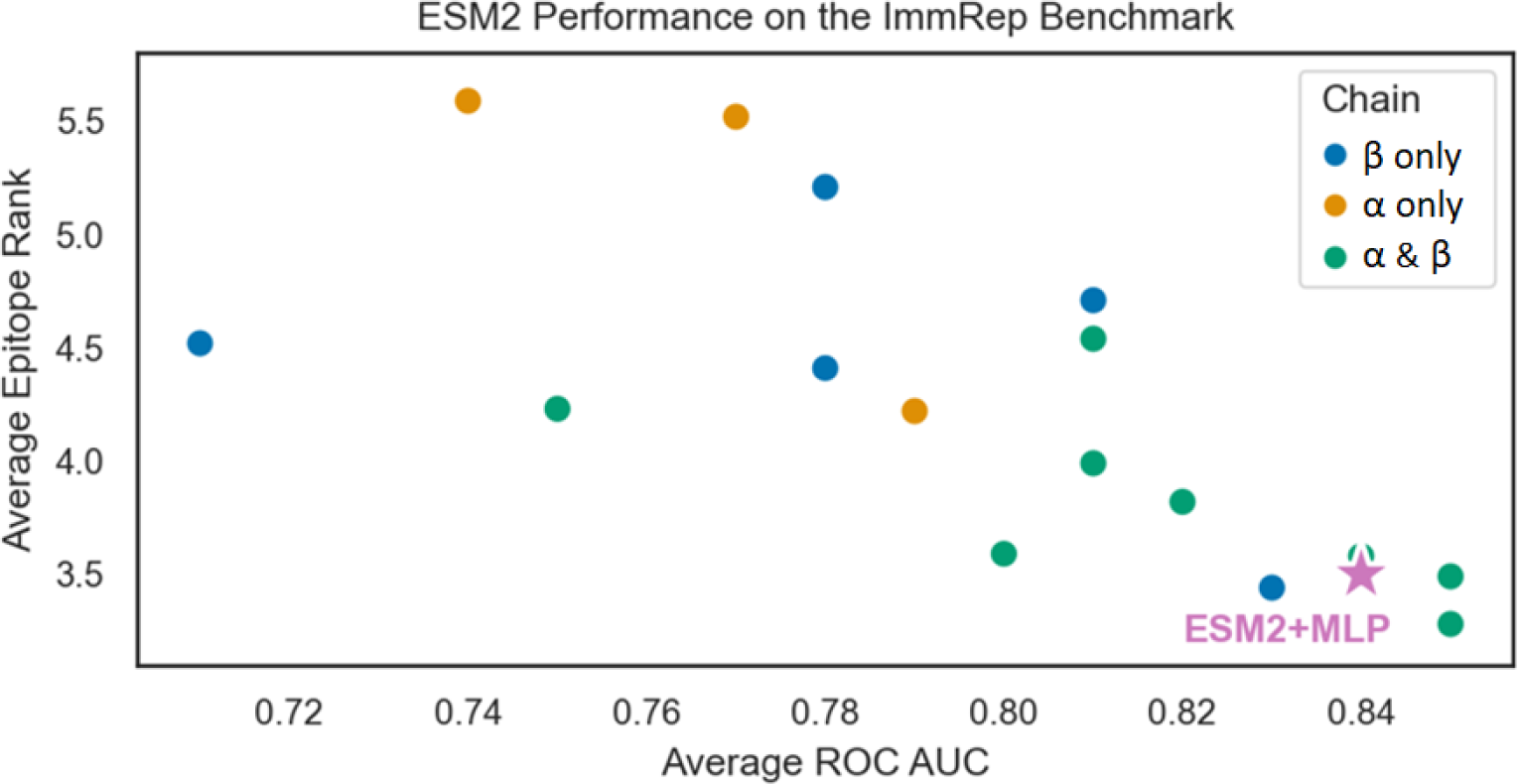
Benchmark performance comparison. Models are compared on the benchmarking test data, on both the epitope ranking (y-axis) and the TCR classification (x-axis) task. Each dot represents one model tested at the ImmRep 2022 benchmark. The pink star indicates the performance of the ESM2-based model, which ranks among the top-performing models in both tasks, located in the bottom right corner. Figure is adapted from [37]

### A.2 Relationship between model size and performance

PLMs are typically large neural networks. In line with the modern understanding of over-parametrized neural networks in the data-rich regime [61, 62, 63], both our own experiments and evidence in the literature [12] tend to indicate that larger PLMs yield better results and more meaningful representation. However, the promise of PLMs as an instantiation of pre-trained foundation models [64] for molecular biology is that downstream application can leverage their learned representation at a much reduced computational cost, and in a lower data regime. Of course, inference with large PLMs is nevertheless computationally expensive, but much less so than training and represents a one-time investment per dataset. As a result, the existence of high-performance small downstream models is desirable to moderate the computational needs of users of PLMs.

**Figure S2:**
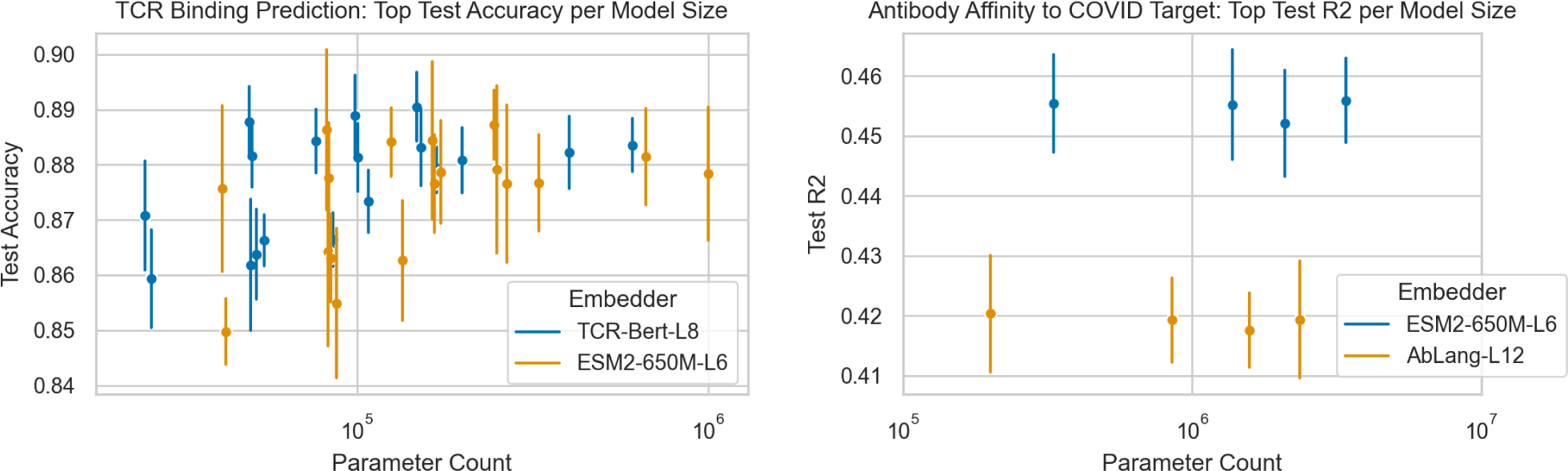
Optimized performance for each tested downstream model size. (Left) for antibody affinity regression. (Right) for TCR binding predictions.

We, therefore, evaluate whether the “larger is better” rule of thumb that works for PLMs also applies to the predictors we trained for antibody affinity regression and TCR binding prediction. To this end, we re-analyze the result of our hyperparameter searches, selecting the best-performing hyperparameter configurations for each total parameter count, based on validation metrics, and observe whether performance was influenced by model size. As we show in Figure S2, there is no visible dependence of optimized performance on parameter count, which is a reassuring signal for bioinformaticians with limited computational resources. Optimized performance metric values can be found in Table S1 and Table S2. Note that the parameters counts we experimentally tested are distributed in a seemingly disordered fashion, which is due to the non-linear relationship between the sampled hidden-layer dimensions and the total number of parameters (Section 4.7).

**Table S1:** Optimal test accuracy for each tested parameter count for the antibody affinity regression task. The reported accuracy is the mean accuracy across replicates and the number in parentheses is the uncertainty on the last significant digit measured as the standard deviation.

| AbLang-L12 |  | ESM-650M-L6 |  |
| --- | --- | --- | --- |
| # Param. | Best Test Acc. | # Param. | Best Test Acc. |
| 200 864 | 0.420(9) | 331 936 | 0.455(8) |
| 852 608 | 0.419(7) | 1 376 896 | 0.455(9) |
| 1 574 912 | 0.417(6) | 2 099 200 | 0.452(8) |
| 2 361 344 | 0.419(9) | 3 409 920 | 0.455(7) |

**Table S2:** Optimal test accuracy for each tested parameter count for the TCR binding prediction task. The reported accuracy is the mean accuracy across replicates and the number in parentheses is the uncertainty on the last significant digit measured as the standard deviation.

| TCR-BERT-L8 |  | ESM-650M-L6 |  |
| --- | --- | --- | --- |
| # Param. | Best Test Acc. | # Param. | Best Test Acc. |
| 24,872 | 0.870(9) | 41,256 | 0.87(1) |
| 25,928 | 0.859(8) | 42,312 | 0.849(5) |
| 49,216 | 0.887(6) | 81,984 | 0.88(1) |
| 49,736 | 0.86(1) | 82,504 | 0.86(1) |
| 50,256 | 0.881(5) | 83,024 | 0.877(9) |
| 51,560 | 0.863(8) | 84,328 | 0.863(7) |
| 54,416 | 0.866(4) | 87,184 | 0.85(1) |
| 76,152 | 0.884(5) | 125,304 | 0.884(6) |
| 85,464 | 0.866(4) | 134,616 | 0.86(1) |
| 98,432 | 0.888(7) | 163,968 | 0.88(1) |
| 100,496 | 0.881(6) | 166,032 | 0.876(8) |
| 107,728 | 0.873(5) | 173,264 | 0.878(9) |
| 147,648 | 0.890(6) | 245,952 | 0.887(6) |
| 152,280 | 0.883(6) | 250,584 | 0.87(1) |
| 168,504 | 0.879(4) | 266,808 | 0.87(1) |
| 198,920 | 0.880(5) | 329,992 | 0.876(8) |
| 401,936 | 0.882(6) | 664,080 | 0.881(8) |
| 609,048 | 0.883(4) | 1,002,264 | 0.87(1) |

## B Supplementary Figures

**Figure S3:**
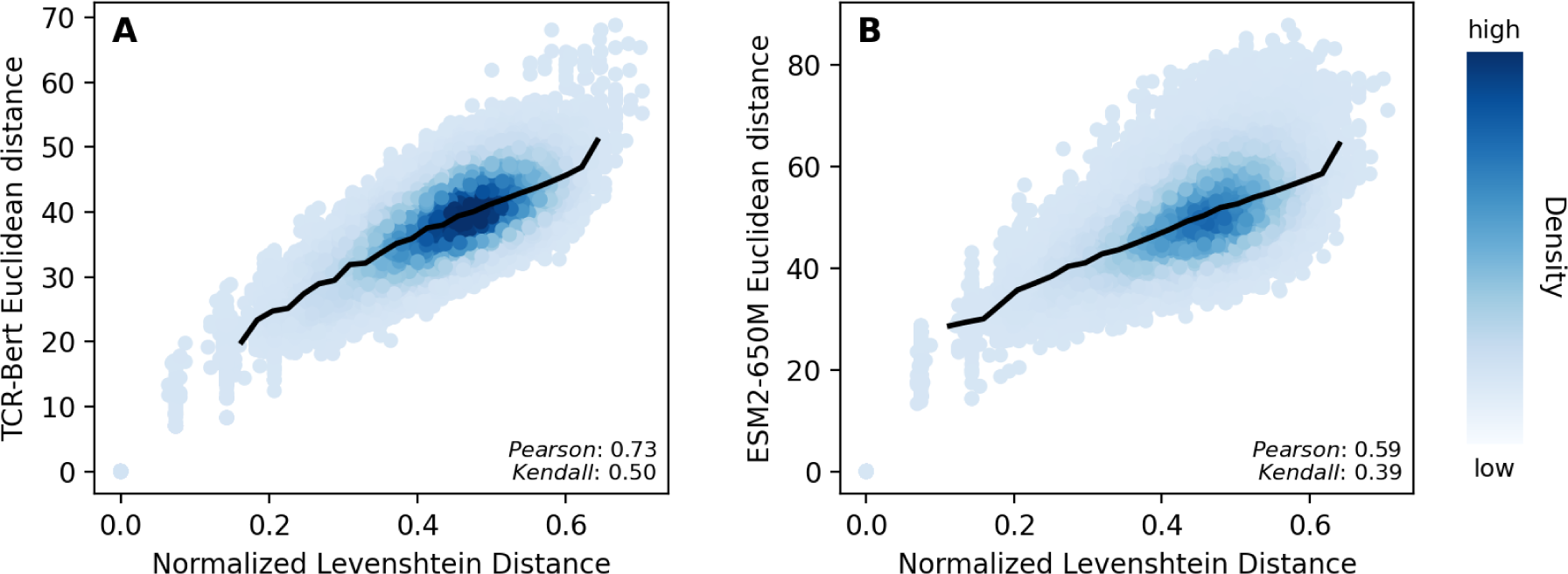
Relationship between Levenshtein and Euclidean distance in the (A) TCR-BERT and (B) ESM2-650M embedding space. The sequences are T Cell receptor CDR3-*β* sequences from [37].

**Figure S4:**
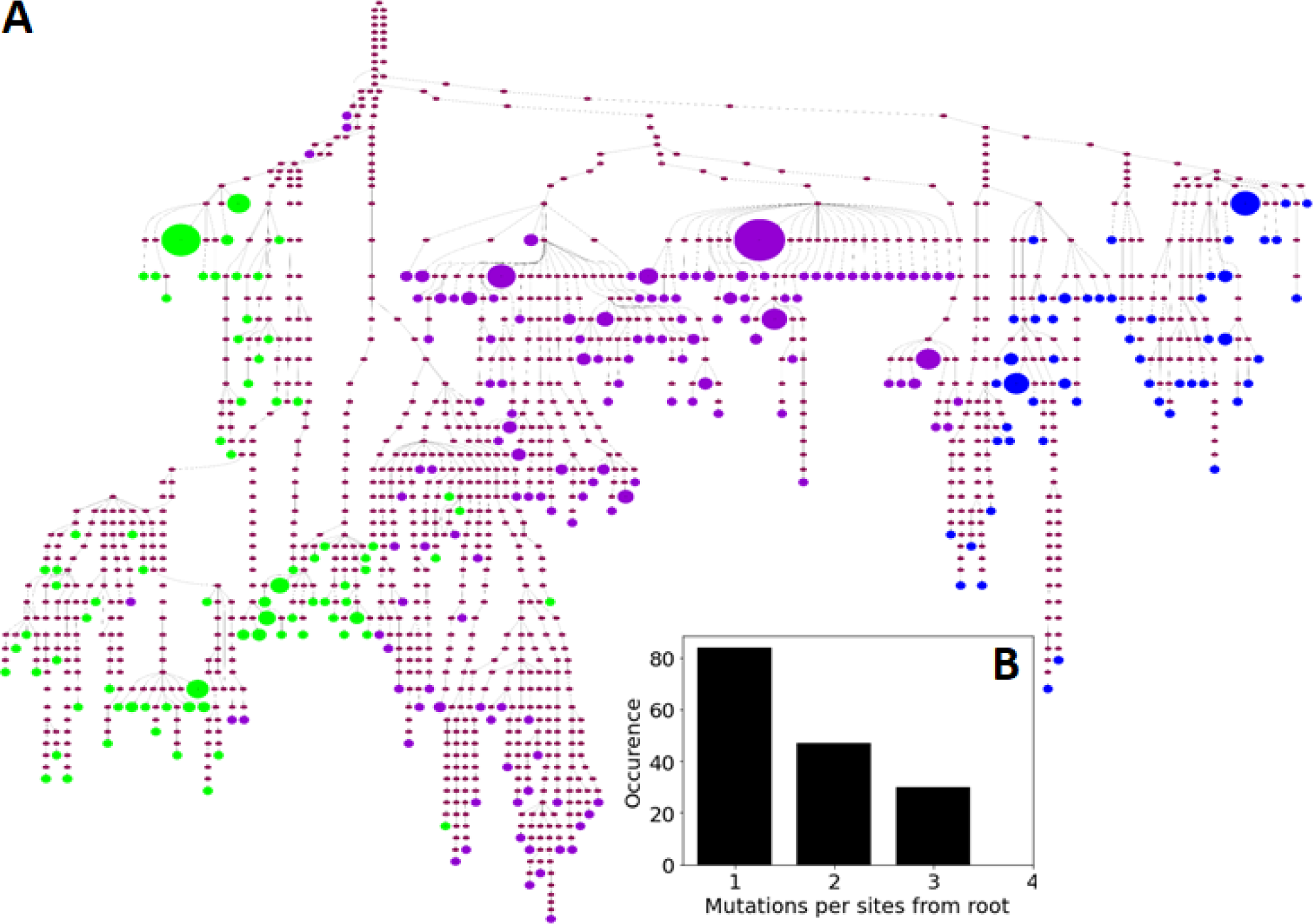
(A) Inferred phylogenetic tree of a B cell clone shared among three GCs. Intermediate brown nodes represent inferred mutations while leaf nodes represent B cells present in the repertoire, colored by their GC. The size of the node indicates the logarithm of the population size for non-unique sequences. We observe up to 300 sequences in the largest node. (B) Number of mutations from the root for each site in the sequence. The infinite site assumption is not verified as more than half of the sites had at least one reoccurring mutation from the root.

**Figure S5:**
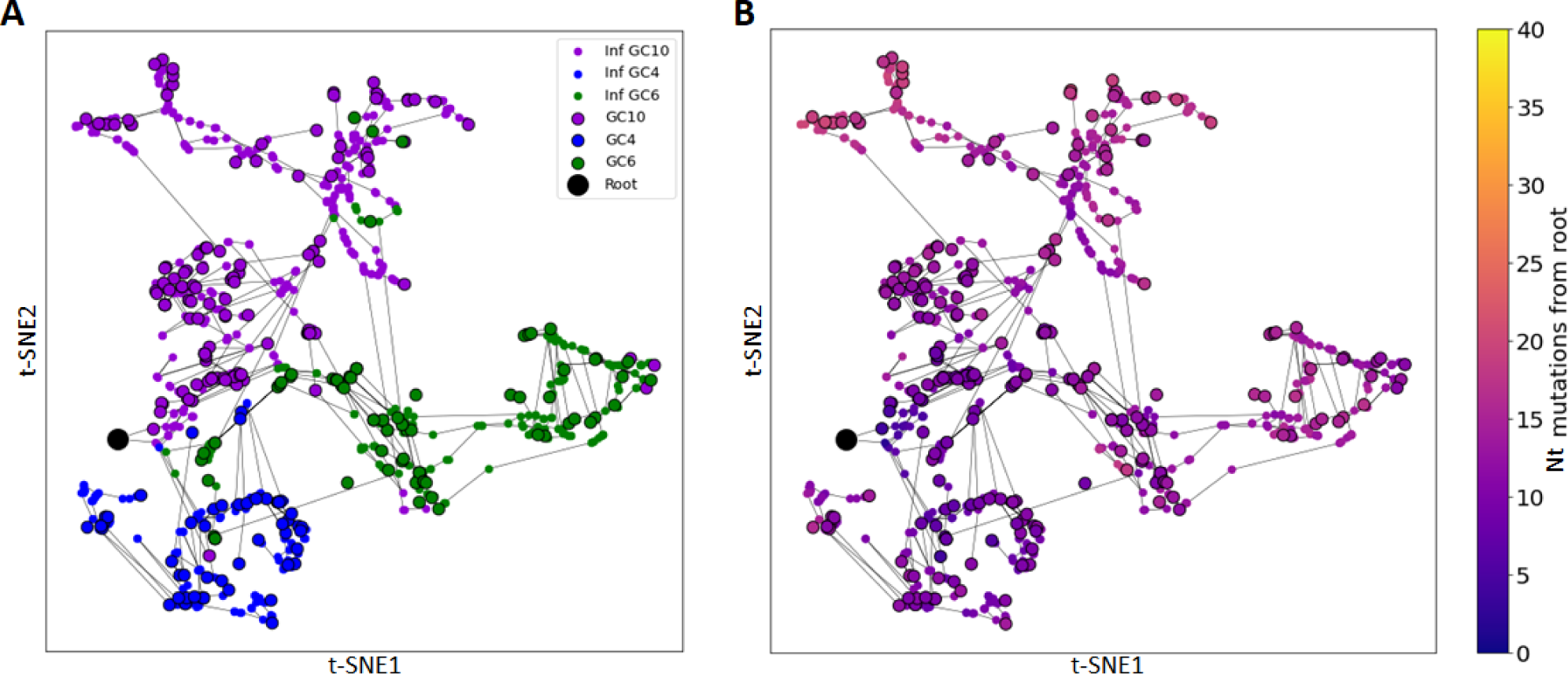
Visualization of phylogenetic relationships using AbLang-embedded BCR sequences. BCRs are colored according to: (A) GC of origin and (B) the number of nucleotide changes from the the root (black dot). Observed sequences are depicted in circled points while inferred intermediate sequences in the phylogeny are in small points and colored according to their nearest neighbor sequence in A. Each line represents a single nucleotide mutation between two sequences.

**Figure S6:**
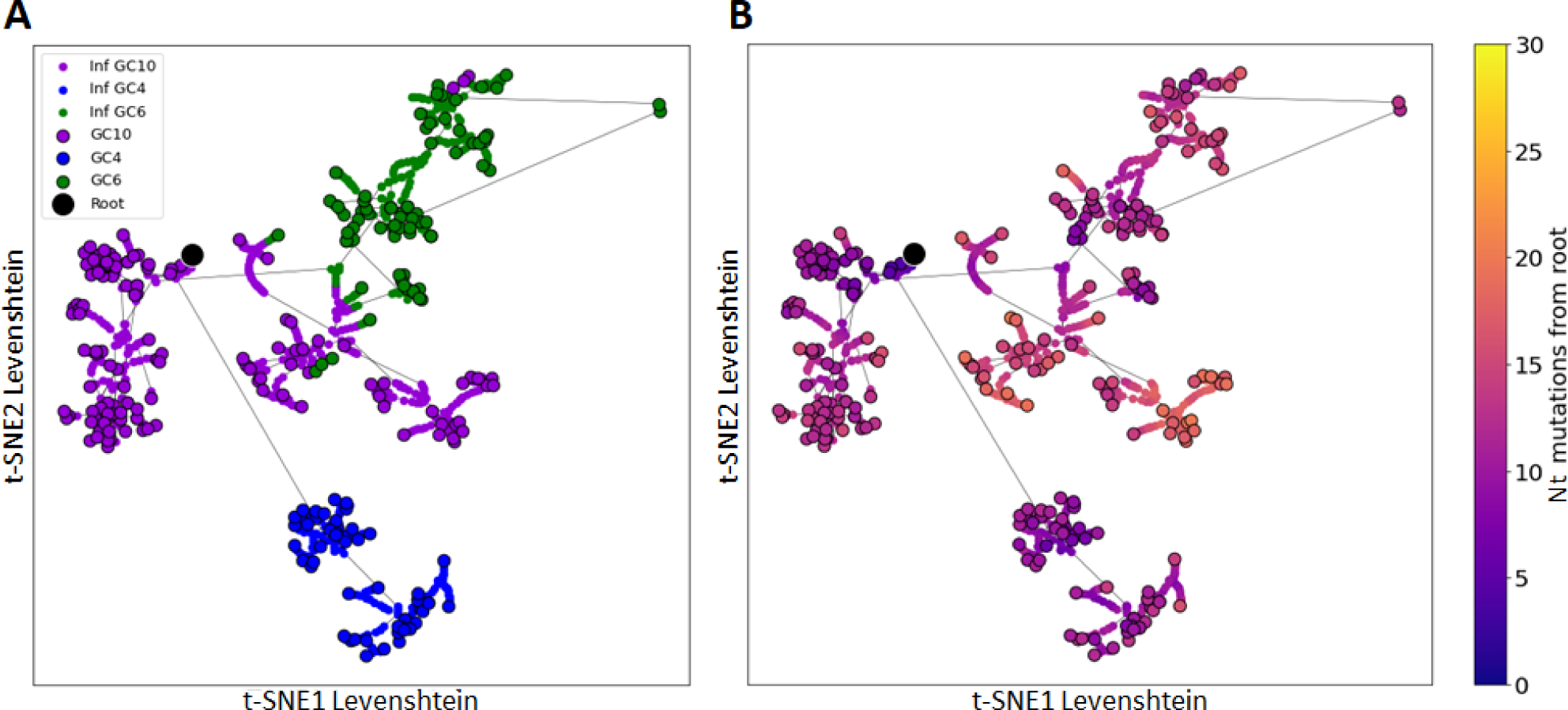
Phylogenetic relationships visualized with a t-SNE representation trained on the pairwise Levenshtein distance between BCR sequences. BCRs are colored according to: (A) GC of origin and (B) the number of nucleotide changes from the the root (black dot). Observed sequences are depicted in circled points while inferred intermediate sequences in the phylogeny are in small points and colored according to their nearest neighbor sequence in A. Each line represents a single nucleotide mutation between two sequences.

**Figure S7:**
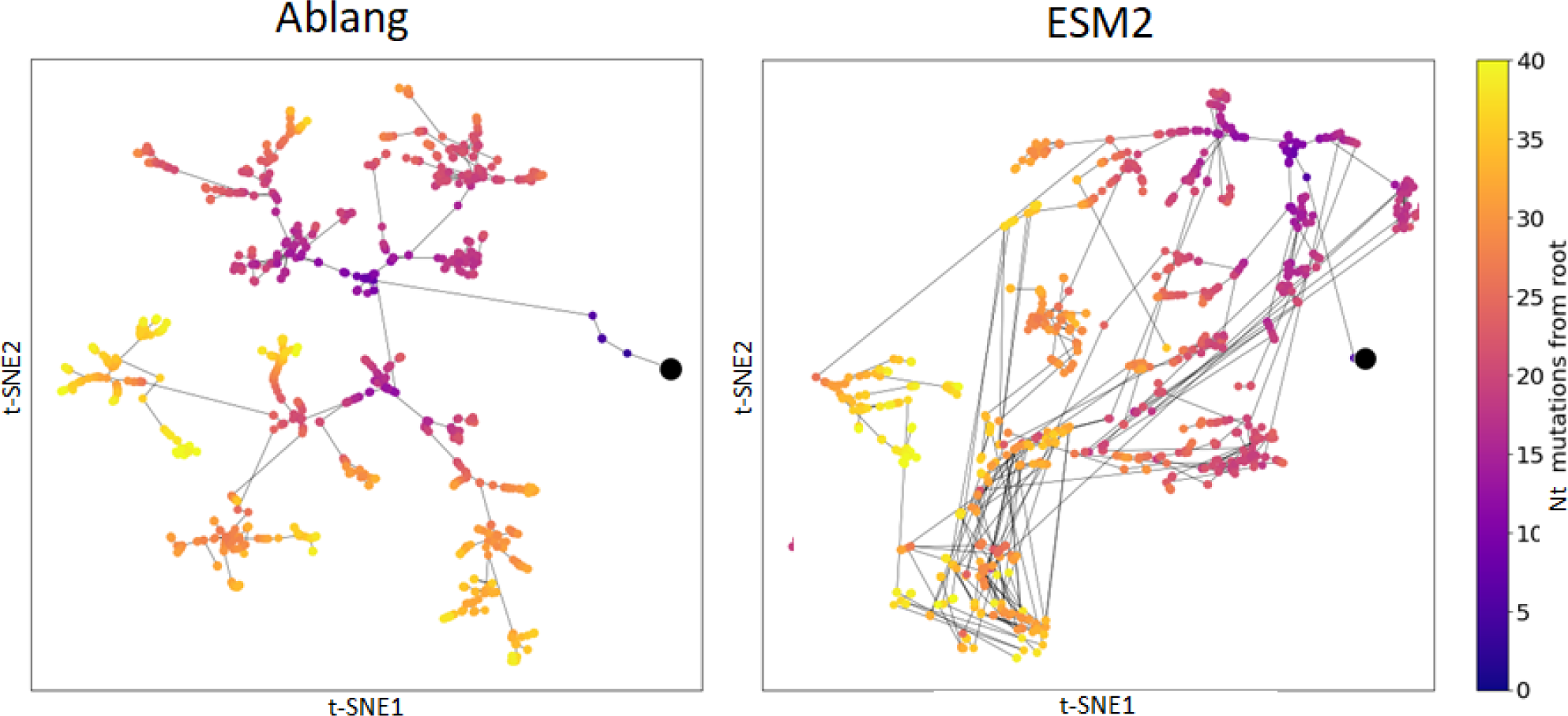
Simulated phylogeny (random mutations with a realistic tree topology) in the AbLang and ESM2-650M embeddings. Each line corresponds to a single nucleotide mutation.

## Notes

### Competing Interest Statement

The authors have declared no competing interest.

### Summary of Updates

spelling mistake in author

## Bibliography

[1] Jacqueline Parkin and Bryony Cohen. “An overview of the immune system”. In: The Lancet 357.9270 (2001), pp. 1777–1789.

[2] Aleksandr Kovaltsuk, et al. “How B-cell receptor repertoire sequencing can be enriched with structural antibody data”. In: Frontiers in immunology 8 (2017), p. 1753.

[3] Rahmad Akbar, et al. “A compact vocabulary of paratope-epitope interactions enables predictability of antibody-antigen binding”. In: Cell Reports 34.11 (2021), p. 108856.

[4] Veronika I Zarnitsyna, et al. “Estimating the diversity, completeness, and cross-reactivity of the T cell repertoire”. In: Frontiers in immunology 4 (2013), p. 485.

[5] Yuval Elhanati, et al. “Inferring processes underlying B-cell repertoire diversity”. In: Philosophical Transactions of the Royal Society B: Biological Sciences 370.1676 (2015), p. 20140243.

[6] Victor Greiff, et al. “Bioinformatic and statistical analysis of adaptive immune repertoires”. In: Trends in immunology 36.11 (2015), pp. 738–749.

[7] Derek M Mason, et al. “Optimization of therapeutic antibodies by predicting antigen specificity from antibody sequence via deep learning”. In: Nature Biomedical Engineering 5.6 (2021), pp. 600–612.

[8] Manasi P Jogalekar, et al. “CAR T-cell-based gene therapy for cancers: new perspectives, challenges, and clinical developments”. In: Frontiers in immunology 13 (2022), p. 925985.

[9] Haig Aghajanian, Joel G Rurik, and Jonathan A Epstein. “CAR-based therapies: opportunities for immuno-medicine beyond cancer”. In: Nature metabolism 4.2 (2022), pp. 163–169.

[10] Tristan Bepler and Bonnie Berger. “Learning the protein language: Evolution, structure, and function”. In: Cell systems 12.6 (2021), pp. 654–669.

[11] Nadav Brandes, et al. “ProteinBERT: a universal deep-learning model of protein sequence and function”. In: Bioinformatics 38.8 (2022), pp. 2102–2110.

[12] Alexander Rives, et al. “Biological structure and function emerge from scaling unsupervised learning to 250 million protein sequences”. In: Proceedings of the National Academy of Sciences 118.15 (2021), e2016239118.

[13] Ratul Chowdhury, et al. “Single-sequence protein structure prediction using a language model and deep learning”. In: Nature Biotechnology 40.11 (2022), pp. 1617–1623.

[14] Ali Madani, et al. “Large language models generate functional protein sequences across diverse families”. In: Nature Biotechnology (2023), pp. 1–8.

[15] Zeming Lin, et al. “Evolutionary-scale prediction of atomic-level protein structure with a language model”. In: Science 379.6637 (2023), pp. 1123–1130.

[16] Tristan Bepler and Bonnie Berger. “Learning protein sequence embeddings using information from structure”. In: International Conference on Learning Representations. 2019.

[17] Zilong Hou, et al. “Learning the protein language of proteome-wide protein-protein binding sites via explainable ensemble deep learning”. In: Communications Biology 6.1 (2023), p. 73.

[18] Wayland Yeung, et al. “Alignment-free estimation of sequence conservation for identifying functional sites using protein sequence embeddings”. In: Briefings in Bioinformatics 24.1 (2023), bbac599.

[19] Tobias H Olsen, Iain H Moal, and Charlotte M Deane. “AbLang: an antibody language model for completing antibody sequences”. In: Bioinformatics Advances 2.1 (2022), vbac046.

[20] Jinwoo Leem, et al. “Deciphering the language of antibodies using self-supervised learning”. In: Patterns 3.7 (2022).

[21] Rohit Singh, et al. “Learning the Language of Antibody Hypervariability”. In: bioRxiv (2023), pp. 2023–04.

[22] Kevin Wu, et al. “TCR-BERT: learning the grammar of T-cell receptors for flexible antigen-xbinding analyses”. In: bioRxiv (2021), pp. 2021–11.

[23] Pengfei Zhang et al. “Context-Aware Amino Acid Embedding Advances Analysis of TCR-Epitope Interactions”. In: (July 2023).

[24] Danqing Wang, YE Fei, and Hao Zhou. “On pre-training language model for antibody”. In: The Eleventh International Conference on Learning Representations. 2023.

[25] Baris E Suzek, et al. “UniRef: comprehensive and non-redundant UniProt reference clusters”. In: Bioinformatics 23.10 (2007), pp. 1282–1288.

[26] Tobias H Olsen, Fergus Boyles, and Charlotte M Deane. “Observed Antibody Space: A diverse database of cleaned, annotated, and translated unpaired and paired antibody sequences”. In: Protein Science 31.1 (2022), pp. 141–146.

[27] Wei Zhang, et al. “PIRD: pan immune repertoire database”. In: Bioinformatics 36.3 (2020), pp. 897– 903.

[28] Mikhail Shugay, et al. “VDJdb: a curated database of T-cell receptor sequences with known antigen specificity”. In: Nucleic acids research 46.D1 (2018), pp. D419–D427.

[29] Vladimir I Levenshtein. “Binary codes capable of correcting deletions, insertions, and reversals”. In: Soviet physics doklady. Vol. 10. 8. Soviet Union. 1966, pp. 707–710.

[30] Aurelien Pelissier, et al. “Exploring the impact of clonal definition on B-cell diversity: implications for the analysis of immune repertoires”. In: Frontiers in Immunology 14 (2023).

[31] Aurelien Pelissier et al. “Convergent evolution and B-cell recirculation in germinal centers in a human lymph node”. In: Life Science Alliance 6.11 (2023). doi: 10.26508/lsa.202301959.

[32] Amos Azaria and Tom Mitchell. “The internal state of an llm knows when its lying”. In: arXiv preprint arXiv:2304.13734 (2023).

[33] Chi Han, et al. “In-Context Learning of Large Language Models Explained as Kernel Regression”. In: arXiv preprint arXiv:2305.12766 (2023).

[34] Kenneth Li, et al. “Emergent world representations: Exploring a sequence model trained on a synthetic task”. In: arXiv preprint arXiv:2210.13382 (2022).

[35] Daniel J Firl, et al. “Capturing change in clonal composition amongst single mouse germinal centers”. In: Elife 7 (2018), e33051.

[36] Victor Greiff, Gur Yaari, and Lindsay G Cowell. “Mining adaptive immune receptor repertoires for biological and clinical information using machine learning”. In: Current Opinion in Systems Biology 24 (2020), pp. 109–119.

[37] Pieter Meysman, et al. “Benchmarking solutions to the T-cell receptor epitope prediction problem: IMMREP22 workshop report”. In: ImmunoInformatics 9 (2023), p. 100024.

[38] Anna Weber, Jannis Born, and Maria Rodriguez Martinez. “TITAN: T-cell receptor specificity prediction with bimodal attention networks”. In: Bioinformatics 37.Supplement_1 (2021), pp. i237– i244.

[39] H. B. Mann and D. R. Whitney. “On a Test of Whether one of Two Random Variables is Stochastically Larger than the Other”. In: The Annals of Mathematical Statistics 18.1 (1947), pp. 50–60. doi: 10.1214/aoms/1177730491. url: 10.1214/aoms/1177730491.

[40] Emily Engelhart, et al. “A dataset comprised of binding interactions for 104,972 antibodies against a SARS-CoV-2 peptide”. In: Scientific Data 9.1 (2022), p. 653.

[41] Umberto Oreste, Alessia Ametrano, and Maria Rosaria Coscia. “On origin and evolution of the antibody molecule”. In: Biology 10.2 (2021), p. 140.

[42] Brian L Hie, et al. “Efficient evolution of human antibodies from general protein language models”. In: Nature Biotechnology (2023).

[43] Aurélien Pélissier, et al. “Computational Model Reveals a Stochastic Mechanism behind Germinal Center Clonal Bursts”. In: Cells 9.6 (2020), p. 1448.

[44] Simone Conti, Edmond Y Lau, and Victor Ovchinnikov. “On the rapid calculation of binding affinities for antigen and antibody design and affinity maturation simulations”. In: Antibodies 11.3 (2022), p. 51.

[45] Rodrigo Garcia-Valiente, et al. “Understanding repertoire sequencing data through a multiscale computational model of the germinal center”. In: npj Systems Biology and Applications 9.1 (2023), p. 8.

[46] Simone Conti, et al. “Multiscale affinity maturation simulations to elicit broadly neutralizing antibodies against HIV”. In: PLoS Computational Biology 18.4 (2022), e1009391.

[47] Jonathan G Faris, et al. “Moving the needle: Employing deep reinforcement learning to push the boundaries of coarse-grained vaccine models”. In: Frontiers in Immunology 13 (2022), p. 1029167.

[48] Li Yujian and Liu Bo. “A normalized Levenshtein distance metric”. In: IEEE transactions on pattern analysis and machine intelligence 29.6 (2007), pp. 1091–1095.

[49] Daniel Mullner. “Modern hierarchical, agglomerative clustering algorithms”. In: arXiv preprint arXiv:1109.2378 (2011).

[50] Namita T Gupta et al. “Hierarchical clustering can identify B cell clones with high confidence in Ig repertoire sequencing data”. In: The Journal of Immunology 198.6 (2017), pp. 2489–2499.

[51] Julie D Thompson, Desmond G Higgins, and Toby J Gibson. “CLUSTAL W: improving the sensitivity of progressive multiple sequence alignment through sequence weighting, position-specific gap penalties and weight matrix choice”. In: Nucleic acids research 22.22 (1994), pp. 4673–4680.

[52] William S DeWitt III, et al. “Using genotype abundance to improve phylogenetic inference”. In: Molecular biology and evolution 35.5 (2018), pp. 1253–1265.

[53] Nika Abdollahi, et al. “Reconstructing B cell lineage trees with minimum spanning tree and genotype abundances”. In: BMC bioinformatics 24.1 (2023), p. 70.

[54] Katharina Jahn, Jack Kuipers, and Niko Beerenwinkel. “Tree inference for single-cell data”. In: Genome biology 17.1 (2016), p. 86.

[55] Yuki Shimoyama. pyCirclize: Circular visualization in Python. Dec. 2022. url: https://github.com/moshi4/pyCirclize.

[56] Adam Paszke et al. “PyTorch: An Imperative Style, High-Performance Deep Learning Library”. In: Advances in Neural Information Processing Systems 32. Curran Associates, Inc., 2019, pp. 8024–8035. url: http://papers.neurips.cc/paper/9015-pytorch-an-imperative-style-high-performance-deep-learning-library.pdf.

[57] Thomas Wolf et al. “Transformers: State-of-the-Art Natural Language Processing”. In: Proceedings of the 2020 Conference on Empirical Methods in Natural Language Processing: System Demonstrations. Online: Association for Computational Linguistics, Oct. 2020, pp. 38–45. url: https://www.aclweb.org/anthology/2020.emnlp-demos.6.

[58] Nils Reimers and Iryna Gurevych. “Sentence-BERT: Sentence Embeddings using Siamese BERT-Networks”. In: Proceedings of the 2019 Conference on Empirical Methods in Natural Language Processing. Association for Computational Linguistics, Nov. 2019. url: https://arxiv.org/abs/1908.10084.

[59] Diederik P. Kingma and Jimmy Ba. “Adam: A Method for Stochastic Optimization”. In: 3rd International Conference on Learning Representations, ICLR 2015, San Diego, CA, USA, May 7-9, 2015, Conference Track Proceedings. Ed. by Yoshua Bengio and Yann LeCun. 2015. url: http://arxiv.org/abs/1412.6980.

[60] Russell D Larsen. “Box-and-whisker plots”. In: Journal of Chemical Education 62.4 (1985), p. 302.

[61] Madhu S. Advani, Andrew M. Saxe, and Haim Sompolinsky. “High-dimensional dynamics of generalization error in neural networks”. In: Neural Networks 132 (2020), pp. 428–446. issn: 0893-6080. doi: 10.1016/j.neunet.2020.08.022. url: https://www.sciencedirect.com/science/article/pii/S0893608020303117.

[62] Mikhail Belkin et al. “Reconciling modern machine-learning practice and the classical bias–variance trade-off”. In: Proceedings of the National Academy of Sciences 116.32 (2019), pp. 15849–15854. doi: 10.1073/pnas.1903070116. eprint: https://www.pnas.org/doi/pdf/10.1073/pnas.19 03070116. url: https://www.pnas.org/doi/abs/10.1073/pnas.1903070116.

[63] Preetum Nakkiran et al. “Deep double descent: Where bigger models and more data hurt”. In: Journal of Statistical Mechanics: Theory and Experiment 2021.12 (2021), p. 124003.

[64] Rishi Bommasani et al. On the Opportunities and Risks of Foundation Models. 2022. arXiv: 2108.0 7258 [cs.LG].

